# Metabolic co-dependence drives the evolutionary ancient *Hydra-Chlorella* symbiosis

**DOI:** 10.1101/234757

**Authors:** Mayuko Hamada, Katja Schröder, Jay Bathia, Ulrich Kürn, Sebastian Fraune, Mariia Khalturina, Konstantin Khalturin, Chuya Shinzato, Nori Satoh, Thomas C.G. Bosch

## Abstract

Many multicellular organisms rely on symbiotic associations for support of metabolic activity, protection, or energy. Understanding the mechanisms involved in controlling such interactions remains a major challenge. In an unbiased approach we identified key players that control the symbiosis between *Hydra viridissima* and its photobiont *Chlorella* sp. A99. We discovered significant upregulation of *Hydra* genes encoding a phosphate transporter and glutamine synthetase suggesting regulated nutrition supply between host and symbionts. Interestingly, supplementing the medium with glutamine temporarily supports in vitro growth of the otherwise obligate symbiotic *Chlorella*, indicating loss of autonomy and dependence on the host. Genome sequencing of *Chlorella* A99 revealed a large number of amino acid transporters and a degenerated nitrate assimilation pathway, presumably as consequence of the adaptation to the host environment. Our observations portray ancient symbiotic interactions as a codependent partnership in which exchange of nutrients appears to be the primary driving force.

## Introduction

Symbiosis has been a prevailing force throughout the evolution of life, driving the diversification of organisms and facilitating rapid adaptation of species to divergent new niches (Moran, 2007; Joy, 2013; McFall-Ngai et al., 2013). In particular, symbiosis with photosynthetic symbiont is observed in many species of Cnidarians such as coral, jellyfish, sea anemone and hydra, contributing to the ecological success of these sessile or planktonic animals (Douglas, 1994; Davy et al., 2012b). Among the many animals dependent on algal symbionts, inter-species interactions between green hydra *Hydra viridissima* and endosymbiotic unicellular green algae of the genus *Chlorella* have been a subject of interest for decades (Muscatine and Lenhoff, 1963; Roffman and Lenhoff, 1969). Such studies not only provide insights into the basic “tool kit” necessary to establish symbiotic interactions, but are also of relevance in understanding the resulting evolutionary selective processes (Muscatine and Lenhoff, 1965a, b; Thorington and Margulis, 1981).

The interactions at play here are clearly metabolic: the algae depend on nutrients that are derived from the host or from the environment surrounding the host, while in return the host receives a significant amount of photosynthetically fixed carbon from the algae. Previous studies have provided evidence that the photosynthetic symbionts provide their host with maltose, enabling *H. viridissima* to survive periods of starvation (Muscatine and Lenhoff, 1963; Muscatine, 1965; Roffman and Lenhoff, 1969; Cook and Kelty, 1982; Huss et al., 1993/1994). *Chlorella*-to-*Hydra* translocation of photosynthates is critical for polyps to grow (Muscatine and Lenhoff, 1965b; Mews, 1980; Douglas and Smith, 1983; Douglas and Smith, 1984). Presence of symbiotic algae also has a profound impact on hydra’s fitness by promoting oogenesis (Habetha et al., 2003; Habetha and Bosch, 2005).

Pioneering studies performed in the 1980s (McAuley and Smith, 1982; Rahat and Reich, 1984) showed that there is a great deal of adaptation and specificity in this symbiotic relationship. All endosymbiotic algae found in a single host polyp are clonal and proliferation of symbiont and host is tightly correlated (Bossert and Dunn, 1986; McAuley, 1986a). Although it is not yet known how *Hydra* controls cell division in symbiotic *Chlorella, Chlorella* strain A99 is unable to grow outside its polyp host and is transmitted vertically to the next generation of *Hydra*, indicating loss of autonomy during establishment of its symbiotic relationship with this host (Muscatine and McAuley, 1982; Campbell, 1990; Habetha et al., 2003).

Molecular phylogenetic analyses suggest that *H. viridissima* is the most basal species in the genus *Hydra* and that symbiosis with *Chlorella* was established in the ancestral *viridissima* group after their divergence from non-symbiotic hydra groups (Martinez et al., 2010; Schwentner and Bosch, 2015). A recent phylogenetic analysis of different strains of green hydra resulted in a phylogenetic tree that is topologically equivalent to that of their symbiotic algae (Kawaida et al., 2013), suggesting these species co-evolved as a result of their symbiotic relationship. Although our understanding of the factors that promote symbiotic relationships in cnidarians has increased (Shinzato et al., 2011; Davy et al., 2012a; Lehnert et al., 2014; Baumgarten et al., 2015; Ishikawa et al., 2016), very little is known about the molecular mechanisms allowing this partnership to persist over millions of years.

Recent advances in transcriptome and genome analysis allowed us to identify the metabolic interactions and genomic evolution involved in achieving the *Hydra-Chlorella* symbiotic relationship. We present here the first characterization, to our knowledge, of genetic complementarity between green *Hydra* and *Chlorella* algae that explains the emergence and/or maintenance of a stable symbiosis. We also provide here the first report of the complete genome sequence from an obligate intracellular *Chlorella* photobiont. Together, our results show that exchange of nutrients is the primary driving force for the symbiosis between *Chlorella* and *Hydra*. Subsequently, reduction of metabolic pathways may have further strengthened their codependency. Our findings provide a framework for understanding the evolution of a highly codependent symbiotic partnership in an early emerging metazoan.

## Results

### Discovery of symbiosis-dependent *Hydra* genes

As tool for our study we used the green hydra *H. viridissima* (**Figure 1A**) colonized with symbiotic *Chlorella* sp. strain A99 (abbreviated here as Hv_Sym), aposymbiotic *H. viridissima* from which the symbiotic *Chlorella* were removed (Hv_Apo), and aposymbiotic *H. viridissima* which had been artificially infected with *Chlorella variabilis* NC64A (Hv_NC64A). The latter is symbiotic to the single-cellular protist Paramecium (Karakashian and Karakashian, 1965). Although an association between *H. viridissima* and *Chlorella* NC64A can be maintained for some time, both their growth rate (**Figure 1B**) and the number of NC64A algae per *Hydra* cell (**Supplementary Figure 1**) is significantly reduced compared to the symbiosis with native symbiotic *Chlorella* A99.

**Figure 1.**
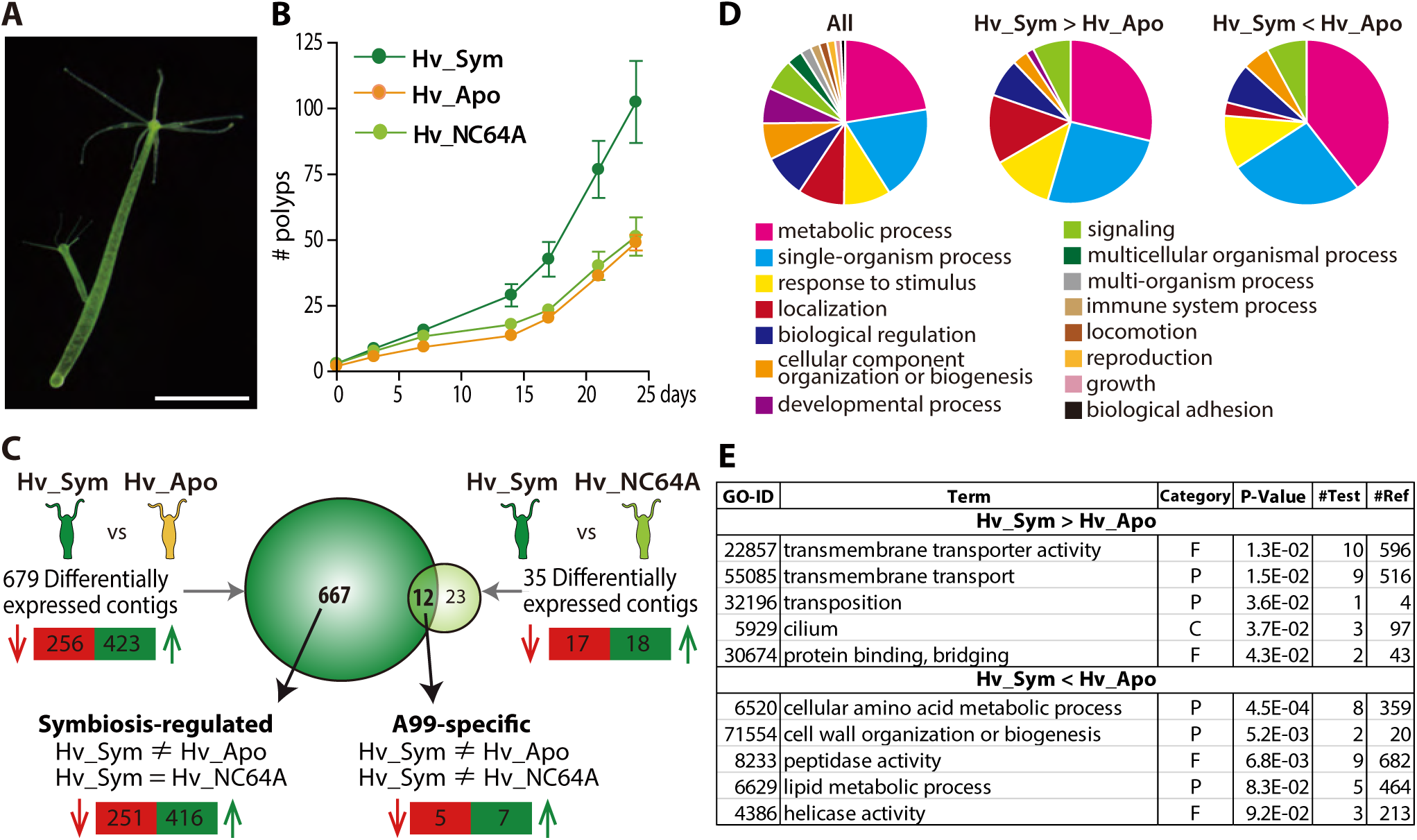
*Hydra* growth and differential expression of Hydra genes resulting from symbiosis. (A) *Hydra viridissima* strain A99 used for this study. Scale bar, 2 mm. (B) Growth rates of polyps grown with native symbiotic Chlorella A99 (Hv_Sym, dark green), Aposymbiotic polyps from which Chlorella were removed (Hv_Apo, orange) and aposymbiotic polyps reinfected with Chlorella variabilis NC64A (Hv_NC64A, light green). (C) Graphic representation of differentially expressed genes identified by microarray. The transcriptome of Hv_Sym is compared with that of Hv_Apo and Hv_NC64A with the number of down-regulated contigs in Hv_Sym shown in red and those up-regulated in green. Genes differentially expressed in Hv_Sym compared to both Hv_Apo and Hv_NC64A are given as “A99-specific”, those differentially expressed between Hv_A99 and Hv_Apo but not Hv_NC64A as “Symbiosis-regulated”. (D) GO distribution of Biological Process at level 2 in all contigs (All), up-regulated contigs (Hv_Sym > Hv_Apo) and down-regulated contigs (Hv_Sym < Hv_Apo) in Hv_Sym. (E) Overrepresented GO terms in up-regulated contigs (Hv_Sym > Hv_Apo) and down-regulated contigs (Hv_Sym < Hv_Apo). Category, F: molecular function, C: cellular component, P: biological process. P-values, probability of Fisher’ s exact test. #Test, number of corresponding contigs in differentially expressed contigs. #Ref, number of corresponding contigs in all contigs.

*H. viridissima* genes involved in the symbiosis with *Chlorella* were identified by microarray based on the contigs of *Hydra viridissima* A99 transcriptome (NCBI GEO Platform ID: GPL23280). For the microarray analysis, total RNA was extracted from the polyps after light exposure for six hours. By comparing the transcriptomes of Hv_Sym and Hv_Apo, we identified 423 contigs that are upregulated and 256 contigs that are downregulated in presence of *Chlorella* A99 (**Figure 1C**). To exclude genes involved in oogenesis and embryogenesis, only contigs differently expressed with similar patterns in both sexual and asexual Hv_Sym were recorded. Interestingly, contigs whose predicted products had no discernible homologs in other organisms including other *Hydra* species were overrepresented in these differentially expressed contigs (Chi-squared test P<0.001) (**Supplementary Figure 2**). Such taxonomically restricted genes (TRGs) are thought to play important roles in the development of evolutionary novelties and morphological diversity within a given taxonomic group (Khalturin et al., 2009; Tautz and Domazet-Loso, 2011).

We further characterized functions of the differentially expressed *Hydra* genes by Gene Ontology (GO) terms (The Gene Ontology et al., 2000). This demonstrated overrepresentation of genes with GO term “localization” in upregulated contigs (Hv_Sym > Hv_Apo) and with GO term “metabolic process” in downregulated contigs (Hv_Sym < Hv_Apo) (**Figure 1D**). More specifically, the upregulated contigs include many genes related to “transmembrane transporter activity”, “transmembrane transport”, “transposition”, “cilium” and “protein binding, bridging” (**Figure 1E**). In the downregulated contig set, the GO classes “cellular amino acid metabolic process”, “cell wall organization or biogenesis” and “peptidase activity” are overrepresented (**Figure 1E**). These results suggest that the *Chlorella* photobiont affects core metabolic processes and pathways in *Hydra*. Particularly, carrier proteins and active membrane transport appears to play a prominent role in the symbiosis.

To narrow down the number of genes specifically affected by the presence of *Chlorella* A99, we identified 12 contigs that are differentially expressed in presence of *Chlorella* A99 but not in presence of *Chlorella* NC64A (**Figure 1C** A99-specific). Independent qPCR confirmed the differential expression pattern for 10 of these genes (**Supplementary Table 1**). The genes upregulated by the presence of the photobiont encode a Spot_14 protein, a glutamine synthetase (GS) and a sodium-dependent phosphate (Na/Pi) transport protein in addition to a *H. viridissima* specific gene (rc_12891: *Sym-1*) and a *Hydra* genus specific gene (rc_13570: *Sym-2*) (**Supplementary Table 1**). *Hydra* genes downregulated by the presence of *Chlorella* A99 were two *H. viridissima* specific genes and three metabolic genes encoding histidine ammonia-lyase, acetoacetyl-CoA synthetase and 2-isopropylmalate synthase (**Supplementary Table 1**). Of the upregulated genes, Spot_14 is described as thyroid hormone-responsive spot 14 protein reported to be induced by dietary carbohydrates and glucose in mammals (Tao and Towle, 1986; Brown et al., 1997). Na/Pi transport protein is a membrane transporter actively transporting phosphate into cells (Murer and Biber, 1996). GS plays an essential role in the metabolism of nitrogen by catalyzing the reaction between glutamate and ammonia to form glutamine (Liaw et al., 1995). Interestingly, out of the three GS genes *H. viridissima* contains only *GS-1* was found to be upregulated by the presence of the photobiont (**Supplementary Figure 3**). The discovery of these transcriptional responses points to an intimate metabolic exchange between the partners in a species-specific manner‥

### Symbiont-dependent *Hydra* genes are upregulated by photosynthetic activity of *Chlorella A99*

To test whether photosynthetic activity of the symbiont is required for upregulation of gene expression, Hv_Sym was either cultured under a standard 12 hr light/dark alternating regime or continuously in the dark for 1 to 4 days prior to RNA extraction (**Figure 2A**). Interestingly, four (*GS1, Spot14, Na/Pi* and *Sym-1*) of five genes specifically activated by the presence of *Chlorella* A99 showed significant upregulation when exposed to light (**Figure 2B**), indicating the relevance of photosynthetic activity of *Chlorella*. This upregulation was strictly dependent on presence of the algae, as in aposymbiont Hv_Apo the response was absent (**Figure 2B**). On the other hand, symbiosis-regulated *Hydra* genes not specific for *Chlorella* A99 (**Figure 1C** Symbiosis-regulated, **Supplementary Table 2**) appear not to be upregulated in a light-dependent manner (**Supplementary Figure 4**). These genes are involved in *Hydra*’s innate immune system (e.g. proteins containing Toll/interleukin-1 receptor domain or Death domain) or in signal transduction (C-type mannose receptor, ephrin receptor, proline-rich transmembrane protein 1, “protein-kinase, interferon-inducible double stranded RNA dependent inhibitor, repressor of (p58 repressor)”). That particular transcriptional changes observed in *Hydra* rely solely on the photosynthetic activity of *Chlorella* A99 was confirmed by substituting the dark incubation with selective chemical photosynthesis inhibitor DCMU (Dichorophenyl-dimethylurea) (Vandermeulen et al., 1972), which resulted in a similar effect (**Figure 2C, D**).

**Figure 2.**
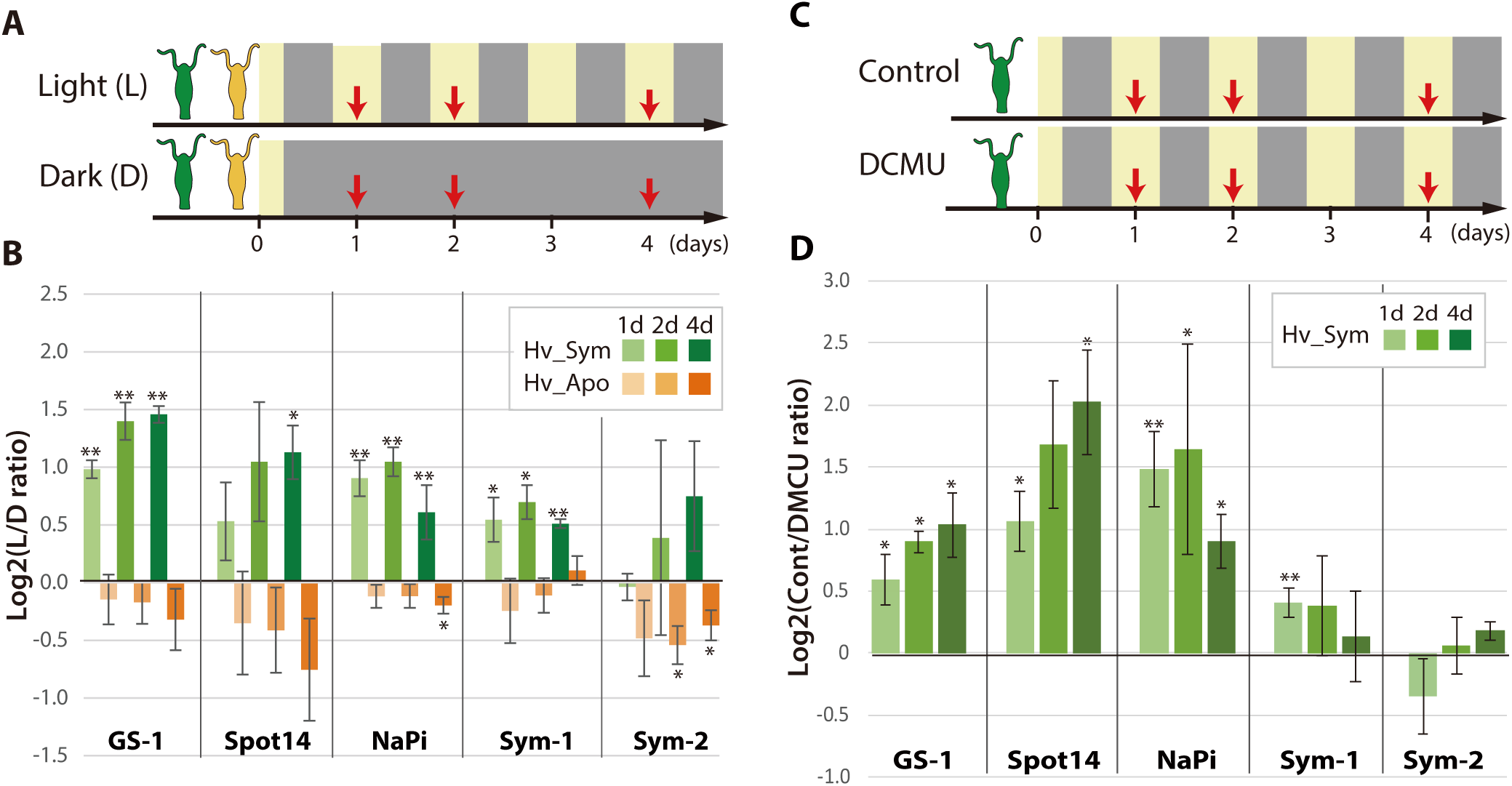
Differential expression of *Hydra* genes under influence of *Chlorella* photosynthesis. (A) Sampling scheme. Hv_Sym (green) and Hv_Apo (orange) were cultured under a standard light-dark regime (Light: L) and in continuous darkness (Dark: D), and RNA was extracted from the polyps at the days indicated by red arrows. (B) Expression difference of five A99-specific genes in Hv_Sym (green bars) and Hv_Apo (orange bars) between the light-dark condition and darkness. The vertical axis shows log scale (log2) fold changes of relative expression level in Light over Dark. (C) Sampling scheme of inhibiting photosynthesis. (D) Differential expression of the five A99-specific genes under conditions allowing (Control) or inhibiting photosynthesis (DCMU). The vertical axis shows log scale (log2) fold changes of relative expression level in Control over DCMU treated. T-tests were performed between Light and Dark

### Symbiont-dependent *Hydra* genes are expressed in endodermal epithelial cells and upregulated by sugars

To further characterize the photobiont induced *Hydra* genes, we performed whole mount *in situ* hybridization (**Figure 3A-F**) and quantified transcripts by qPCR using templates from isolated endoderm and ectoderm (**Supplementary Figure 5**), again comparing symbiotic and aposymbiotic polyps (**Figure 3 G-I**). The GS-1 gene and the Spot14 gene are expressed both in ectoderm and in endoderm (**Figure 3A, B**) and both genes are strongly upregulated in the presence of the photobiont (**Figure 3G, H**). In contrast, the Na/Pi gene was expressed only in the endoderm (**Figure 3C**) and there it was strongly upregulated by the photobiont (**Figure 3I**). Since *Chlorella* sp. A99 colonizes endodermal epithelial cells only, the impact of algae on symbiosis-dependent genes in both the ectodermal and the endodermal layer indicates that photosynthetic products can be transported across these two tissue layers or some signals can be transduced by cell-cell communication.

**Figure 3.**
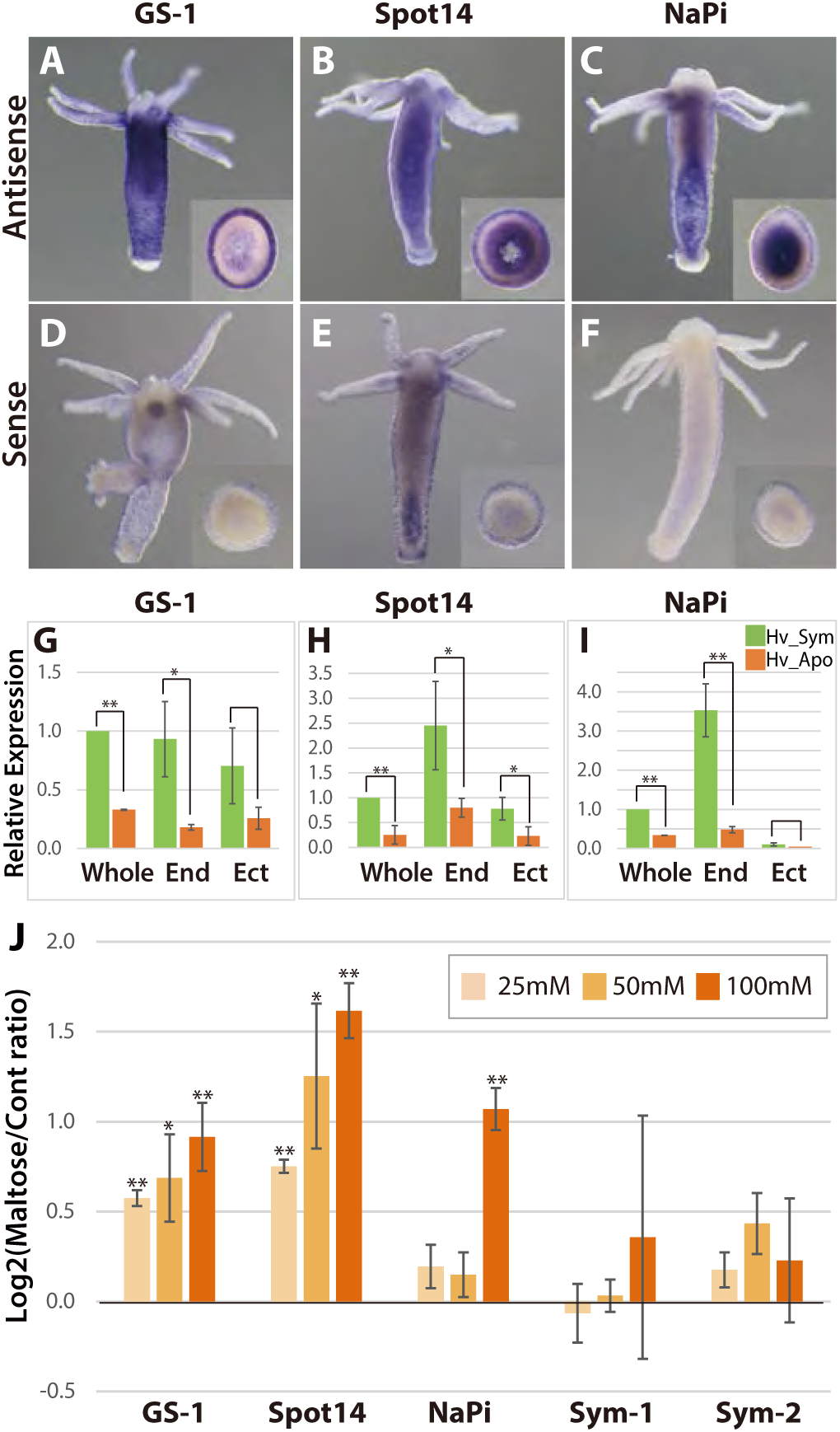
Spatial expression patterns of genes coding for glutamine synthetase, Spot 14 and Na/Pi-transporter. A-F; Whole mount in situ hybridization using antisense (A-C) and sense probes (D-F; negative controls) for glutamine synthetase-1 (GS-1; left), Spot 14 (center) and Na/Pi-transporter (NaPi; right). Inserts show cross sections of the polyp’ s body. (G-I) Relative expression levels of whole animal (whole), isolated endoderm (End) and isolated ectoderm (Ect) tissue of Hv_Sym (green bars) and Hv_Apo (orange bars). T-test was performed between Hv_Sym and Hv_apo. Pvalue, * <0.05, ** <0.01. (J) Expression change of genes GS-1, Spot14, NaPi, Sym-1 and Sym-2 following exposure to 25mM, 50mM and 100mM maltose in Hv_Apo. The vertical axis shows log scale (log2) fold changes of relative expression level of maltose-treated over the untreated Hv_Apo control. T-test was performed between maltose-treated and control. Pvalue, * <0.05, ** <0.01.

To more closely dissect the nature of the functional interaction between *Hydra* and *Chlorella* and to explore the possibility that maltose released from the algae is involved in A99-specific gene regulation, we cultured aposymbiotic polyps (Hv_Apo) for 2 days in medium containing various concentrations of maltose (**Figure 3J**). Of the five A99 specific genes, GS-1 and the Spot14 gene were upregulated by maltose in a dose-dependent manner; the Na/Pi gene was only upregulated in 100mM maltose and the *Hydra* specific genes Sym-1 and Sym-2 did not show significant changes in expression by exposure to maltose (**Figure 3J**). This provides strong support for previous views that maltose excretion by symbiotic algae contributes to the stabilization of this symbiotic association (Cernichiari et al., 1969). When polyps were exposed to glucose instead of maltose, the genes of interest were also transcriptionally activated in a dose-dependent manner, while sucrose had no effect (**Supplementary Figure 6A-D**). Exposure to low concentrations of galactose increased transcriptional activity but at high concentration it did not, indicating a substrate inhibitor effect for this sugar. That the response to glucose is similar or even higher compared to maltose after 6 hours of treatment (**Supplementary Figure 6E**), suggests that *Hydra* cells transform maltose to glucose as a source of energy.

### The *Chlorella A99* genome records a symbiotic life style

To better understand the symbiosis between *H. viridissima* and *Chlorella* and to refine our knowledge of the functions that are required in this symbiosis, we sequenced the genome of *Chlorella* sp. strain A99 and compared it to the genomes of other green algae. The genome of *Chlorella* sp. A99 was sequenced to approximately 211-fold coverage, enabling the generation of an assembly comprising a total of 40.9 Mbp (82 scaffolds, N50=1.7Mbp) (**Supplementary Table 3**). *Chlorella* sp. A99 belongs to the family *Chlorellaceae* (**Figure 4A**) and of the green algae whose genomes have been sequenced it is most closely related to *Chlorella variabilis* NC64A (NC64A) (Merchant et al., 2007; Palenik et al., 2007; Worden et al., 2009; Blanc et al., 2010; Prochnik et al., 2010; Blanc et al., 2012; Gao et al., 2014; Pombert et al., 2014). The genome size of the total assembly in strain A99 was similar to that of strain NC64A (46.2Mb) (**Figure 4B**). By k-mer analysis (k-mer = 19), the genome size of A99 was estimated to be 61 Mbp (Marcais and Kingsford, 2011). Its GC content of 68%, is the highest among the green algae species recorded (**Figure 4B**). In the A99 genome, 8298 gene models were predicted. As shown in **Figure 4C**, about 80% of these predicted genes have extensive sequence similarity to plant genes, while 13% so far have no similarity to genes of any other organisms (**Figure 4C**). It is also noteworthy that 7% of the A99 genes are similar to genes of other kingdoms but not to *Hydra*, indicating the absence of gene transfer from *Hydra* to the photobiont genome (**Figure 4C**).

**Figure 4.**
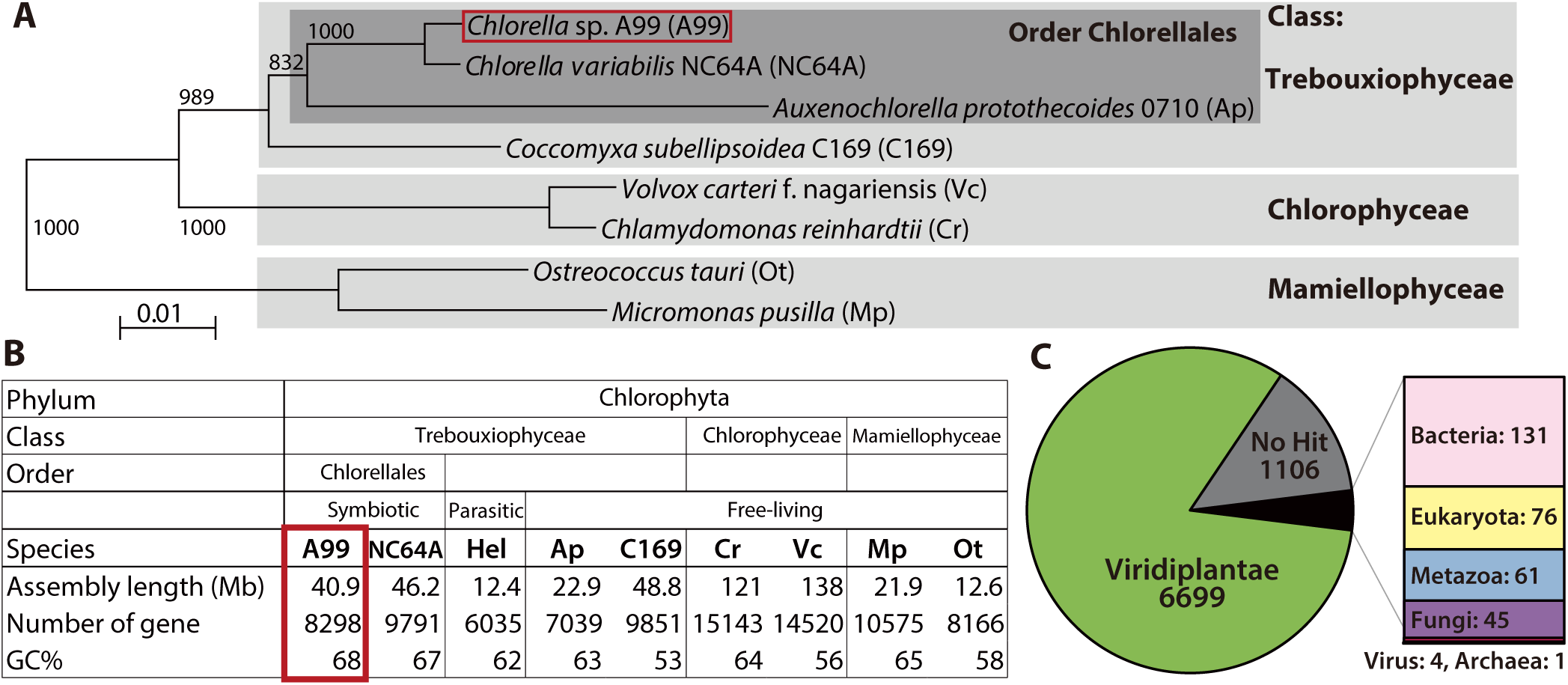
Comparison of key features deduced from the *Chlorella* A99 genome with other green algae. (A) Phylogenetic tree of eight genome sequenced chlorophyte green algae including ***Chlorella*** sp. A99. The NJ tree is based on sequences of the 18S rRNA gene, ITS1, 5.8S rRNA gene, ITS2 and 28S rRNA gene. (B) Genomic features and taxonomy of the sequenced chlorophyte green algae. Hel: *Helicosporidium* sp. ATCC50920. (C) The proportion of similarity of Chlorella A99 gene models to those of other organisms.

### The *Chlorella* A99 genome provides evidences for extensive nitrogenous amino acid import and an incomplete nitrate assimilation pathway

Several independent lines of evidence demonstrate that nitrogen limitation and amino-acid metabolism have a key role in the *Chlorella–Hydra* symbiosis and that symbiotic *Chlorella* A99 depends on glutamine provided by its host (Rees, 1986; McAuley, 1987a, b, 1991; Rees, 1991) (Rees, 1989). To identify *Chlorella* candidate factors for the development and maintenance of the symbiotic life style, we therefore used the available genome information to assess genes potentially involved in amino acid transport and the nitrogen metabolic pathway.

When performing a search for the Pfam domain “Aa_trans” or “AA_permease” to find amino acid transporter genes in the A99 genome, we discovered numerous genes containing the Aa_trans domain (**Supplementary Table 4A**). In particular, A99 contains many orthologous genes of amino acid permease 2 and of transmembrane amino acid transporter family protein (solute carrier family 38, sodium-coupled neutral amino acid transporter), as well as NC64A (**Supplementary Table 4B, C**). Both of these gene products are known to transport neutral amino acids including glutamine. This observation is supporting the view that import of amino acids is an essential feature for the symbiotic way of life of *Chlorella*.

In nitrogen assimilation processes, plants usually take up nitrogen in the form of nitrate (NO_3_^-^) via nitrate transporters (NRTs) or as ammonium (NH_4_^+^) via ammonium transporters (AMT) (**Figure 5A**). In higher plants, two types of nitrate transporters, NRT1 and NRT2, have been identified (Krapp et al., 2014). Some NRT2 require nitrate assimilation-related component 2 (NAR2) to be functional (Quesada et al., 1994). NO_3_^-^ is reduced to nitrite by nitrate reductase (NR), NO_2_^-^ is transported to the chloroplast by nitrate assimilation-related component1 (NAR1), and NO_2_^-^ is reduced to NH_4_^+^ by nitrite reductase (NiR). NH_4_^+^ is incorporated into glutamine (Gln) by glutamine synthetase (GS), and Gln is incorporated into glutamate (Glu) by NADH-dependent glutamine amide-2-oxoglutarate aminotransferase (GOGAT), also known as glutamate synthase. This pathway is highly conserved among plants. In the genomes of 10 green algae species sequenced so far, the major components of the pathway, including NRT1 and NRT2, NAR1 and NAR2, NR, NiR, AMT, GOGAT and GS, are all present, although NRT1 is absent in the *Micromonas pusilla* genome (Sanz-Luque et al., 2015).

**Figure 5.**
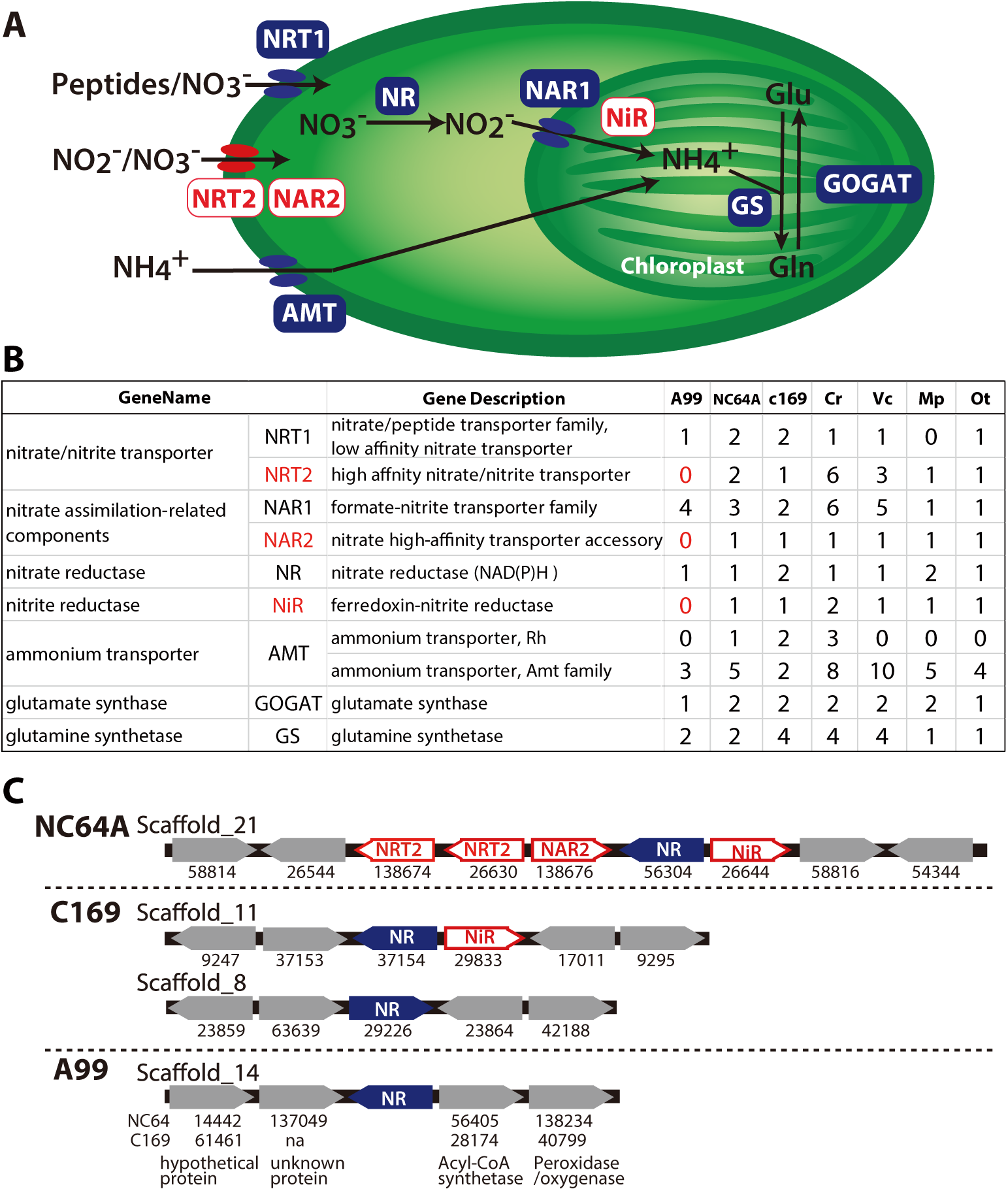
Nitrogen assimilation pathways in *Chlorella* A99. (A) Schematic diagram of the nitrogen assimilation pathway in plants showing the function of nitrate transporters NRT1 (peptides/nitrate transporter) and NRT2 (nitrate/nitrite transporter), nitrate assimilation-related components NAR1 and NAR2, nitrate reductase NR, nitrite reductase NiR, ammonium transporter AMT, glutamate synthetase GOGAT and glutamine synthetase GS. Genes shown in red boxes (NRT2, NAR2 and NiR) were not found in the *Chlorella* sp. A99 genome. (B) Table showing the number of nitrogen assimilation genes in *Chlorella* sp. A99 (A99), *Chlorella variabilis* NC64A (NC64A), *Coccomyxa subellipsoidea* C169 (C169), *Volvox carteri* f. nagariensis (Vc), *Chlamydomonas reinhardtii* (Cr), *Ostreococcus tauri* (Ot) and *Micromonas pusilla* (Mp). (C) Gene clusters of nitrate assimilation genes around the shared NR genes (blue) in the genomes of NC64A, C169 and A99. Red boxes show nitrate assimilation genes absent in A99 and gray boxes depict other genes. Numbers below the boxes are JGI protein IDs of NC64A and C169. Numbers below the genes of A99 are JGI protein IDs of the best hit genes in NC64A and C169 and their gene name.

Based on the annotation by Sanz-Luque et al. (Sanz-Luque et al., 2015), we searched these nitrogen assimilation genes in the *Chlorella* A99 genome, using ortholog grouping and a reciprocal blast search using the protein sequences from other green algae (**Figure 5B**, **Supplementary Table 5**). As expected, the *Chlorella* A99 genome contains many homologues of the genes involved in nitrogen assimilation in plants including genes encoding NRT1, NAR1, NR, AMT, GS and GOGAT (**Figure 5B**). Intriguingly, our systematic searches have failed to identify representative genes for NRT2, NAR2 and NiR in the *Chlorella* A99 genome (**Figure 5B**). We confirmed the absence of the NRT2 and NiR genes by PCR using primers designed for the conserved regions of these genes and which failed to produce a product with genomic DNA as a template (**Supplementary Figure 7**). Due to the weak sequence conservation of the NAR2 gene in the three algae genomes, PCR of that gene was not performed. Taken together, our observations indicate that *Chlorella* A99 algae appear to lack NRT2, NAR2 and NiR.

Since in many fungi, cyanobacteria and algae species, nitrate assimilation genes are known to act in concert and a gene cluster of NR and NiR genes is conserved between different green algae (Sanz-Luque et al., 2015), we next investigated the level of genomic clustering of the nitrate assimilation pathway genes in the *Chlorella* genome. Comparing the genomes of NC64A and *Coccomyxa subellipsoidea* C169 (C169) revealed the presence of a cluster of NR and NiR genes (**Figure 5C**). In NC64A, two NRT2 genes, together with genes for NAR2, NR and NiR are clustered on scaffold 21. In C169, one of NR genes and NiR are clustered together but the second NR gene is separate. Interestingly, analyzing the sequences around the NR gene in the *Chlorella* A99 genome provided no evidence for the presence of a co-localized NiR gene or any other nitrate assimilation genes, nor any conserved gene synteny to NC64A and C169 (**Figure 5C**). Our comparative genomic analyses therefore points to an incomplete as well as scattered nitrogen metabolic pathway in symbiotic *Chlorella* A99, which lacks essential transporters and enzymes for nitrate assimilation and also lacks the clustered structure of nitrate assimilation genes.

### Supplementing the medium with glutamine allows temporary *in vitro* growth of symbiotic *Chlorella* A99

The absence of genes essential for nitrate assimilation in the *Chlorella* A99 genome (**Figure 5**) is consistent with its inability to grow outside the *Hydra* host cell (Habetha and Bosch, 2005) and indicates that *Chlorella* symbionts are dependent on metabolites provided by their host. We hypothesized that *Chlorella* is unable to use nitrite and ammonium as a nitrogen source, and that it relies on *Hydra* assimilating ammonium to glutamine to serve as the nitrogen source. To test this hypothesis and to examine utilization of nitrogen compounds of A99, we isolated *Chlorella* A99 from Hv_Sym and cultivated it *in vitro* using modified bold basal medium (BBM) (Nichols and Bold, 1965) containing the same amount of nitrogen in the form of NO_3_^-^, NH_4_^+^, Gln or casamino acids (**Figure 6**, **Supplementary Table 6**). As controls, *Chlorella variabilis* NC64A (NC64A) isolated from Hv_NC64A and free-living C169 were used. To confirm that the cultured A99 is not contamination, we amplified and sequenced the genomic region of the 18S rRNA gene by PCR (**Supplementary Figure 8**) and checked this against the genomic sequence of A99. Kamako et al. reported that free-living algae *Chlorella vulgaris* Beijerinck var. *vulgaris* grow in media containing only inorganic nitrogen compounds as well as in media containing casamino acids as a nitrogen source, while NC64A required amino acids for growth (Kamako et al., 2005). Consistent with these observations, C169 grew in all tested media and NC64A grew in media containing casamino acids and Gln, although its growth rate was quite low in presence of NH_4_^+^ and NO_3_^-^ (**Figure 6**). Remarkably, *Chlorella* A99 increased in cell number for up to 8 days in media containing casamino acids and Gln (**Figure 6**). Similar to NC64A, A99 did not grow in presence of NH_4_^+^ and NO_3_^-^. The growth rates of both A99 and NC64A were higher in medium containing a mixture of amino acids (casamino acids) than the single amino acid Gln. In contrast to NC64A, A99 could not be cultivated permanently in casamino acids or glutamine supplemented medium, indicating that additional growth factors are necessary to maintain *in vitro* growth of this obligate symbiont. Thus, although *in vitro* growth of A99 can be promoted by adding Glu and amino acids to the medium, A99 cannot be cultured permanently in this enriched medium, indicating that other host derived factors remain to be uncovered.

**Figure 6.**
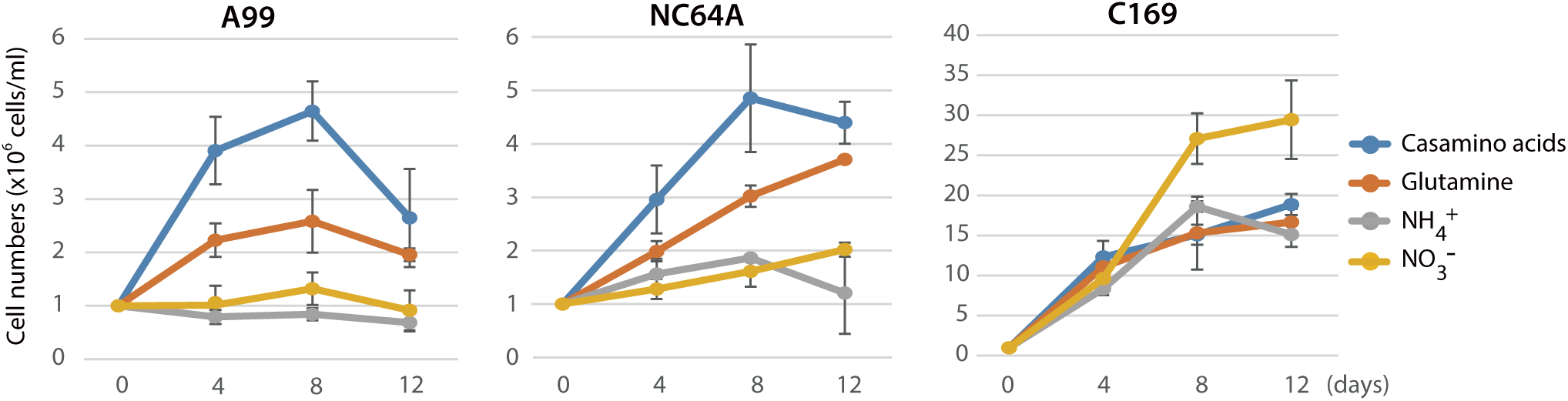
Growth of green algae in presence of various nitrogen sources. The growth rate of *Chlorella* A99 (A99), *Chlorella variabilis* NC64A (NC64A) and *Coccomyxa subellipsoidea* C-169 (C169) by in vitro culture was assessed for different nitrogen sources with casamino acids (blue), glutamine (orange), ammonium (gray) and nitrate (yellow). Mean number of algae per ml were determined at 4, 8, 12 days after inoculation with 106 cell/ml.

## Discussion

Sequencing of the *Chlorella* A99 genome in combination with the transcriptome analyses of symbiotic, aposymbiotic and NC64A-infected *H. viridissima* polyps has enabled the identification of genes with specific functions in this symbiotic partnership. The *Hydra-Chlorella* symbiosis links carbohydrate supply from the photobiont to glutamine synthesis by the host. Characteristics of the symbiont genome obviously reflect its adaptation to this way of life, including an increase in amino acid transporters and degeneration of the nitrate assimilation pathway. This conclusion is based on six observations: **(i)** Expression of some genes including GS-1, Spot 14 and NaPi is specifically upregulated in the presence of *Chlorella* A99 (**Fig. 1C**, **Supplementary Table 1**), and **(ii)** they are induced by both, photosynthetic activity of *Chlorella* and by supplying exogenous maltose or glucose (**Figure 2**, **3J**, **Supplementary Figure 6**). These results indicate that maltose release by photosynthesis of the symbiont enhances nutrition supply including glutamine by the host (**Figure 7**). **(iii)** Symbiotic *Chlorella* A99 cannot be cultivated *in vitro* in medium containing a single inorganic nitrogen source (**Figure 6**). Since medium containing glutamine supports *in vitro* growth of A99, this organism appears to depend on glutamine provided by the *Hydra* host. **(i*v*)** The genome of *Chlorella* A99 contains multiple amino acid transporter genes (**Supplementary Table 4**), but lacks genes involved in nitrate assimilation (**Figure 5**), pointing to amino acids as main source of nitrogen and a degenerated nitrate assimilation pathway. As for ammonium, which is one of the main nitrogen sources in plants, previous studies have reported the inability of symbiotic algae to take up ammonium because of the low peri-algal pH (pH 4-5) that stimulates maltose release (Douglas and Smith, 1984; Rees, 1989; McAuley, 1991; Dorling et al., 1997). Since *Chlorella* apparently cannot use nitrite and ammonium as a nitrogen source, it seems that *Hydra* has to assimilate ammonium to glutamine and provides it to *Chlorella* A99 (**Figure 7**).

**Figure. 7.**
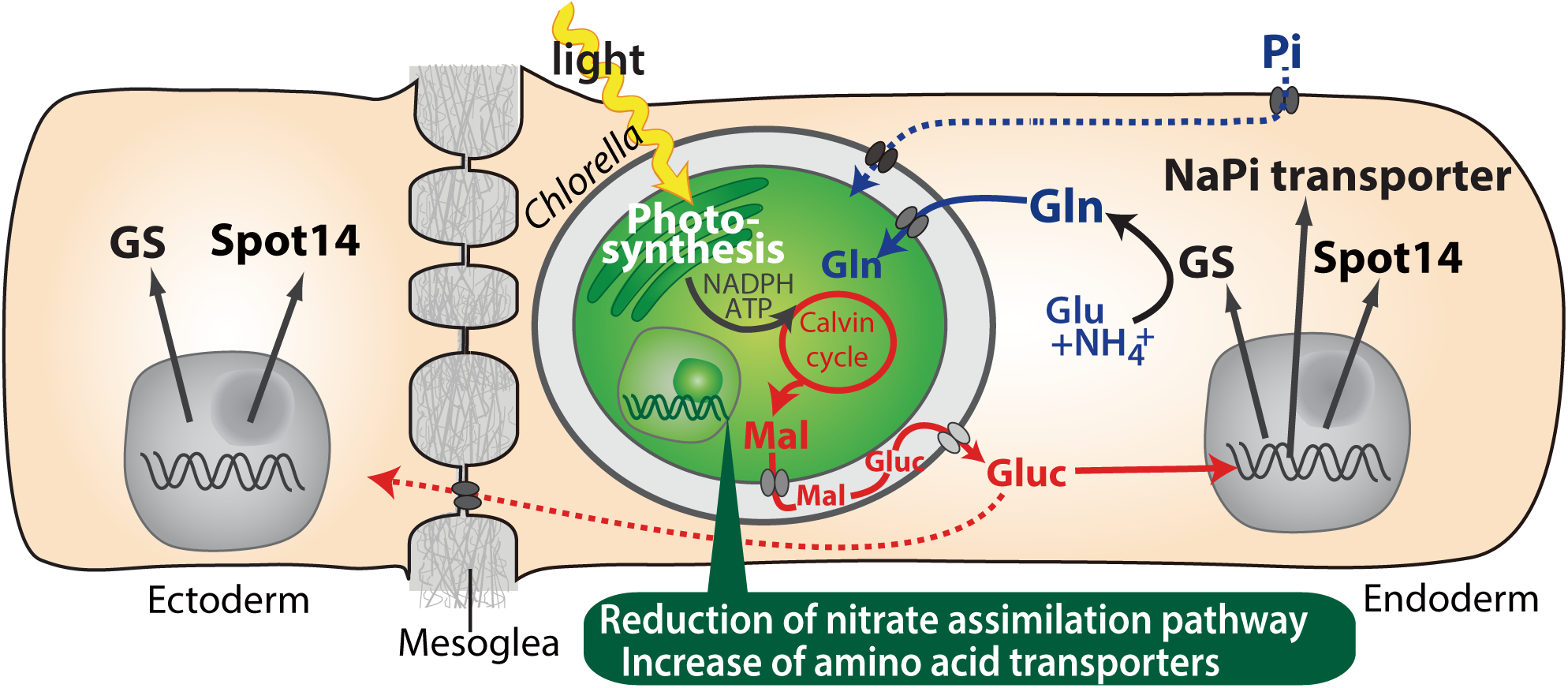
Molecular interactions in the symbiosis between green hydra and Chlorella A99. During light conditions, *Chlorella* performs photosynthesis and produces maltose (Mal) which is secreted into the Hydra symbiosome where it is possibly digested to glucose (Gluc), shown in red. The sugar induces expression of hydra genes encoding glutamine synthetase (GS), Na/Pi transporter (NaPi) and Spot14. GS synthesizes glutamine (Gln) from glutamate (Glu) and ammonium (NH4+). Gln is used by *Chlorella* as a nitrogen source. Since the sugar also upregulates the gene for NaPi which controls intracellular phosphate levels, it might be involved in the supply of phosphorus to *Chlorella* as well (blue broken line). The sugar is transmitted or defused to the ectoderm (red broken line) and there induces the expression of GS and Spot14. In the *Chlorella* A99 genome, degeneration of the nitrate assimilation system and an increase of amino acid transporters was observed (green balloon).

**(*v*)** While polyps with native symbiont *Chlorella* A99 grew faster than aposymbiotic ones, symbiosis with foreign algae NC64A had no effect on the growth of polyps at all (**Figure 1B**). **(*vi*)** *Hydra* endodermal epithelial cells host significantly fewer NC64A algae than A99 (**Supplementary Figure 1**) providing additional support for the view of a tightly regulated codependent partnership in which exchange of nutrients appears to be the primary driving force.

Previous studies have reported that symbiotic *Chlorella* in green hydra releases significantly larger amounts of maltose than NC64A (Mews and Smith, 1982; Rees, 1989). In addition, Rees reported that *Hydra* polyps containing high maltose releasing algae had a high GS activity, whereas aposymbiotic *Hydra* or *Hydra* with a low maltose releasing algae had lower GS activity (Rees, 1986). Although the underlying mechanism of how maltose secretion and transportation from *Chlorella* is regulated is still unclear, the amount of maltose released by the symbiont could be an important symbiont-derived driver or stabilizer of the *Hydra–Chlorella* symbiosis.

Exchange of nitrogenous compounds and photosynthetic products between host and symbiont is widely found in other symbiotic associations. For example, in marine invertebrates such as corals, sea anemones, and giant clams associated with *Symbiodinium* algae, the algae provide the photosynthate in forms of glucose, glycerol, organic acids, amino acids or lipids to their host, and in turn the symbionts receive ammonia or glutamine as nitrogen sources (Burriesci et al., 2012; Davy et al., 2012; Kellogg and Patton, 1983; Lewis and Smith, 1971; Muscatine, 1965; Muscatine and Cernichiari, 1969; 1993; Trench, 1971; Venn et al., 2008; Whitehead and Douglas, 2003; Yellowlees et al., 2008). Moreover, in corals a Na/Pi transporter is involved in the uptake of phosphate across host membranes, and the zooxanthellae contribute to the uptake of phosphate (D’Elia, 1977; Jackson et al., 1989). These observations together with the results presented here make the host-controlled supply of nitrogen and phosphorus as a response of a signal photosynthate seem the universal principle of invertebrate-algae symbiosis.

Metabolic dependence of symbionts on host supply occasionally results in genome reduction and gene loss. For example, the symbiotic *Buchnera* bacteria of insects are missing particular genes in essential amino acid pathways (Shigenobu et al., 2000; Hansen et al., 2011). The fact that the corresponding genes of the host are upregulated in the bacteriocyte, indicates complementarity and syntrophy between host and symbiont. Similarly, in *Chlorella* A99 the nitrogen assimilation system could have been lost as result of continuous supply of nitrogenous amino acids provided by *Hydra*. On the other hand, the genome size and total gene number of *Chlorella* A99 is similar to other species in the class Trebouxiophyceae (**Figure 4B**). The apparently unchanged complexity of the Chlorella A99 genome suggests a relatively early stage of this symbiotic partnership. From these observation, we propose that the gene loss in metabolic pathway is the first step of genome reduction caused by dependency on nutrients from the host. Our study suggests metabolic-codependency is the primary driving force for the evolution of symbiosis between *Hydra* and *Chlorella*.

## Materials and methods

### Biological materials and procedures

Experiments were carried out with the Australian *Hydra viridissima* strain A99, which was obtained from Dr. Richard Campbell, Irvine. Polyps were maintained at 18°C on a 12 hours light/dark cycle and fed with *Artemia* two or three times a week. Aposymbiotic (algae free) polyps were obtained by photobleaching using 5 μM DCMU (3-(3,4-dichlorophenyl)-1,1-dimethylurea) as described before (Pardy, 1976; Habetha et al., 2003). Experiments were carried out with polyps starved for 3-6 days. Isolation of endodermal layer and ectodermal layer was performed as described by Kishimoto et al. (Kishimoto et al., 1996). Symbiotic *Chlorella* were isolated as described before by Muscatine and McAuley (Muscatine, 1983; McAuley, 1986b). *Chlorella variabilis* NC64A (NIES-2541), *Coccomyxa subellipsoidea* C-169 (NIES-2166) and *Chlamydomonas reinhardtii* (NIES-2235) were obtained from the Microbial Culture Collection at the National Institute for Environmental Studies (Tsukuba, Japan).

### Nucleic acid preparation

Total RNA of *Hydra* was extracted by use of the Trizol reagent and PureLink RNA Mini Kit (Life Technology) after lysis and removal of algae by centrifugation. The genomic DNA of green algae was extracted using ISOPLANT II (Nippon Gene, Tokyo, Japan) following DNase I treatment to degrade contaminant DNA. Quantity and quality of DNA and RNA were checked by NanoDrop (Thermo Scientific Inc., Madison, USA) and BioAnalyzer (Agilent Technologies, Santa Clara, USA).

### Microarray Analysis

cRNA targets labeled with cyanine-3 were synthesized from 400 ng total *Hydra* RNA using a Quick Amp Labeling Kit for one color detection (Agilent Technologies). A set of fluorescently labeled cRNA targets was employed in a hybridization reaction with 4 × 44K Custom-Made *Hydra viridissima* Microarray (Agilent Technologies) contributing a total of 43,222 transcripts that was built by mRNA-seq data (NCBI GEO Platform ID: GPL23280) (Bosch et al., 2009).

Hybridization and washing were performed using the GE Hybridization Kit and GE Wash Pack (Agilent Technologies) after which the arrays were scanned on an Agilent Technologies G2565BA microarray scanner system with SureScan technology following protocols according to the manufacturer’s instructions. The intensity of probes was extracted from scanned microarray images using Feature Extraction 10.7 software (Agilent Technologies). All algorithms and parameters used in this analysis were used with default conditions. Background-subtracted signal-intensity values (gProcessedSignal) generated by the Feature Extraction software were normalized using the 75^th^ percentile signal intensity among the microarray. Those genes differentially expressed between two samples were determined by average of fold change (cut off >2.0) and Student’s t-test (P< 0.1). The data series are accessible at NCBI GEO under accession number GSE97633.

### Quantitative real time RT-PCR

Total RNA was reverse transcribed using First Strand cDNA Synthesis Kit (Fermentas, Ontario, Canada). Real-time PCR was performed using GoTaq qPCR Master Mix (Promega, Madison, USA) and ABI Prism 7,300 (Applied Biosystems, Foster City, USA). All qPCR experiments were performed in duplicate with three biological replicates each. Values were normalized using the expression of the tubulin alpha gene. Primers used for these experiments are listed in **Supplementary Table 7A**.

### Whole mount *in situ* hybridization

Expression patterns of specific *Hydra* genes were detected by whole mount *in situ* hybridization with digoxigenin (DIG)-labelled RNA probes. Specimens were fixed in 4% paraformaldehyde. Hybridization signal was visualized using anti-DIG antibodies conjugated to alkaline phosphatase and NBT/BCIP staining solution (Roche). DIG-labeled sense probes (targeting the same sequences as the antisense probes) were used as a control. Primers used for these experiments are listed in **Supplementary Table 7B**.

### Genome sequencing and gene prediction

For genome sequencing of *Chlorella* sp. A99, *Chlorella* sp. A99 was isolated from *H. viridissima* A99 and genomic DNA was extracted. Paired-end library (insert size: 740 bp) and mate-pair libraries (insert size: 2.2 and 15.2kb) were made using Illumina TruSeq DNA LT Sample Prep Kit and Nextera Mate Pair Sample Preparation Kit respectively (Illumina Inc., San Diego, USA), following the manufacturer’s protocols. Genome sequencing was performed using Illumina Miseq and Hiseq 2000 platforms. Sequence reads were assembled using Newbler Assembler version 2.8 (Roche, Penzberg, Germany) and subsequent scaffolding was performed by SSPACE (Boetzer et al., 2011). Gaps inside the scaffolds were closed with the paired-end and mate-pair data using GapCloser of Short Oligonucleotide Analysis Package (Luo et al., 2012). To overcome potential assembly errors arising from tandem repeats, sequences that aligned to another sequence by more than 50% of the length using blastn (1e-50) were removed from the assembly. The completeness of the genome was evaluated using CEGMA v2.4 (Core Eukaryotic Genes Mapping Approach) based on mapping of the 248 most highly conserved core eukaryotic genes (CEGs) on the assembled genome (Parra et al., 2007). The completeness of complete and partial CEGs in the A99 scaffolds was 80% and 88%, respectively. The fraction of repetitive sequences was 12%. Gene models was predicted by AUGUSTUS 3.0.1 using model parameters for NC64A (Stanke et al., 2006). This Whole Genome Shotgun project has been deposited at DDBJ/ENA/GenBank under the accession PCFQ00000000 (BioProject ID: PRJNA412448). The genome sequences and gene models are also accessible at website of OIST Marine Genomics Unit Genome Project (http://marinegenomics.oist.jp/chlorellaA99/viewer/info?project_id=65).

### Analysis of genes in *Hydra viridissima* and *Chlorella*

Annotation of transcriptome contigs and prediction of gene models was performed by use of BLAST, Gene Ontology (The Gene Ontology et al., 2000) and blast2go (Conesa et al., 2005). To examine the conservation of *H. viridissima* contigs among metazoans, homology searches by blastx (evalue 1E-5) were performed using protein databases obtained from NCBI for *Drosophila melanogaster* and *Homo sapiens*, from the JGI genome portal (http://genome.jgi.doe.gov/) for *Branchiostoma floridae, Nematostella vectensis*, from Echinobase (http://www.echinobase.org/EchinoBase/) for *Strongylocentrotus pupuratus*, from Compagen for *Hydra magnipapillata*, and from the OIST marine genomics Genome browser ver.1.1 (http://marinegenomics.oist.jp/coral/viewer/info?project_id=3) for *Acropora digitifera*.

For comparative analysis of gene models of *Chlorella* sp. A99 and other algae, domain searches against the Pfam database (Pfam-A.hmm) were performed using HMMER (Eddy, 1998; Finn et al., 2016), and ortholog gene grouping was done using OrthoFinder (Emms and Kelly, 2015). The sequences of the reference genes and genomes were obtained from the database of the JGI genome portal for *Chlorella variabilis* NC64A, *Coccomyxa subellipsoidea* C-169, *Volvox carteri, Micromonas pusilla*, and *Ostreococcus tauri*, from NCBI for *Auxenochlorella protothecoides* 0710 and from Phytozome (http://phytozome.jgi.doe.gov/pz/portal.html) for *Chlamydomonas reinhardtii* (Merchant et al., 2007; Worden et al., 2009; Blanc et al., 2010; Prochnik et al., 2010; Blanc et al., 2012; Gao et al., 2014; Pombert et al., 2014)

Nitrogen assimilation genes in *Chlorella* A99 were identified by orthologous gene groups and reciprocal blast searches. The number of genes for nitrate assimilation genes, glutamine synthetase and glutamate synthetase, and clustering of such genes were systematically reported by (Sanz-Luque et al., 2015). We used these data as reference for searches of nitrogen assimilation genes, and further nitrogen assimilation genes were searched by Kyoto Encyclopedia of Genes and Genomes (KEGG) pathway (Kanehisa and Goto, 2000). JGI genome browsers of *Chlorella variabilis* NC64A and *Coccomyxa subellipsoidea* C-169 were also used for retrieving genes and checking gene order on the scaffolds.

### Phylogenetic analysis

For a phylogenetic tree of chlorophyte green algae, the sequences of 18S rRNA gene, ITS1, 5.8S rRNA gene, ITS2 and 28S rRNA gene were obtained from scaffold20 of *Chlorella* A99 genome sequence, and from NCBI nucleotide database entries for *Chlorella variabilis* NC64A (FM205849.1), *Auxenochlorella protothecoides* 0710 (NW_011934479.1), *Coccomyxa subellipsoidea* C169 (AGSI01000011.1), *Volvox carteri* f. nagariensis (NW_003307662.1), *Chlamydomonas reinhardtii* (FR865576.1), *Ostreococcus tauri* (GQ426340.1) and *Micromonas pusilla* (FN562452.1). Multiple alignments were produced with CLUSTALX (2.1) with gap trimming (Larkin et al., 2007). Sequences of poor quality that did not well align were deleted using BioEdit (Hall, 1999). Phylogenetic analyses were performed using the Neighbor-Joining method by CLUSTALX. Representative phylogenetic trees were drawn by using NJ plot (Perriere and Gouy, 1996).

### PCR amplification of nitrate assimilation genes in green algae

Primers were designed based on the conserved region of the NRT2 gene, NiR and NR genes (positive control) identified by comparison of genes from *Chlorella variabilis* NC64A (NC64A), *Coccomyxa subellipsoidea* C169 (C169), and *Chlamydomonas reinhardtii* (Cr) which belongs to Chlorophyceae class of green algae. Primers for NAR2 could not be designed because of insufficient conservation. As positive controls, amplicons were produced for NR of all the green algae examined and of NRT2 and NiR from NC64A, C169 and Cr, after which their sequences were checked. KOD FX Neo (TOYOBO, Tokyo, Japan) was used under the following conditions: an initial denaturation phase (94 °C for 120 sec) followed by 36 cycles of (98 °C for 30 sec, 69 °C for 100 sec) for NiR, (98 °C for 30 sec, 58 °C for 30 sec and 68 °C for 210 sec) for NRT2 and (98 °C for 30 sec, 59 °C for 30 sec and 68 °C for 60 sec) for NR. In each case, 10 ng gDNA was used as a template. The primers used are described in **Supplementary Table 7C**. PCR products were sequenced to confirm amplification of the target genes using ABI PRISM 3100 Genetic Analyzer (Thermo Fisher Scientific Inc., Madison, USA) using BigDye Terminator v3.1 Cycle Sequencing Kit (Thermo Fisher Scientific).

### *In vitro* culture of algae

To isolate symbiotic algae, polyps were quickly homogenized in 0.25% sodium dodecyl sulfate (SDS) solution and centrifuged at 3000g for 1 min. The pellet was resuspended in 0.05% SDS and centrifuged at 500g for 5min. Isolated A99, NC64A and C169 were washed by sterilized Bold Basal Medium (Bischoff and Bold, 1963) modified by the addition of 0.5% glucose, 1.2mg/L vitamine B1 (Thiaminhydrochloride), 0.01mg/L vitamine B12 (Cyanocobalamin) (**Supplementary Table 6**) and incubated for two days in modified Bold Basal Medium with 50mg/l ampicillin and streptomycin. The algae were cultivated in 5 ml of modified Bold Basal Medium (BBM) with the same amount of nitrogen (2.9 mM NaNO_3_, NH_4_Cl, glutamine or 426 mg/l casamino acids) and 5mg/l Carbendazim (anti-fungal) with fluorescent illumination (12 hour light, 12 hour dark) at 20°C. Mean numbers of algae per ml were calculated from three tubes enumerated at 4, 8, and 12 days after inoculation with 10^6^ cell/sml using a hemocytometer. After cultivation, gDNA was isolated from the A99 cultured in Gln-containing BBM and casamino acid-containing BBM and A99 was isolated from green hydra directly. A partial genomic region of the 18S rRNA gene was amplified by PCR and sequenced to confirm absence of contamination by other algae. PCR was performed using AmpliTaq Gold (Thermo Fisher Scientific). Sequencing was performed as described above. The primers used are described in **Supplementary Table 7D**.

## Acknowledgement

We thank Trudy Wassenaar for critical reading the text and for discussion. We also thank the DNA Sequencing Section in OIST for genome sequencing. This work was supported in part by JSPS KAKENHI Grant-in-Aid for Young Scientists (B) 25840132 and Scientific Research (C) 15K07173 to M.H. and by the Deutsche Forschungsgemeinschaft (DFG) (CRC1182 “Origin and Function of Metaorganisms”). T.C.G.B. gratefully appreciates support from the Canadian Institute for Advanced Research (CIFAR).

## Competing interests

The authors declare that no competing interests exist.

## Supplemental Tables

**Supplemental Table 1.** List of the A99 specific genes differentially expressed in Hv_Sym compared to both Hv_Apo and Hv_NC64A and fold changes of expression level examined by microarray and qPCR.

A99>Apo, NC64A
| Probe name (gene ID) | Microarray |  | qPCR |  | Seq Description (1) | InterProScan |
| --- | --- | --- | --- | --- | --- | --- |
|  | A99/Apo | A99/NC64A | A99/Apo | A99/NC64A |  |  |
| rc_13579 | 12.8 | 4.0 | 11.2 | 4.0 | uncharacterized protein LOC101238438 [Hydra vulgaris] |  |
| rc_12891 | 9.0 | 2.9 | 14.6 | 6.9 | (Hydra viridis specific) |  |
| 27417 | 4.5 | 4.8 | 3.0 | 3.0 |  | IPR009786 Spot_14 |
| rc_26218 | 3.3 | 2.4 | 2.5 | 2.3 |  | PTHR10010 Sodium-dependent phosphate transport protein 2C |
| 1046 | 3.1 | 2.1 | 2.2 | 1.6 | glutamine synthetase-like |  |

A99<Apo, NC64A
| Probe name (gene ID) | Microarray |  | qPCR |  | Seq Description | InterProScan |
| --- | --- | --- | --- | --- | --- | --- |
|  | Apo/A99 | NC64A/A99 | Apo/A99 | NC64A/A99 |  |  |
| NPNHRC_26859 | 83.2 | 9.7 | ∞ | ∞ | (Hydra viridis specific) | |
| RC_FVQRUGK01AXSJ | 13.7 | 2.6 | 2.1 | 1.5 | acetoacetyl-CoA synthetase-like |  |
| rc_14793 | 7.2 | 4.1 | 9.4 | 4.8 | 2-isopropylmalate synthase | IPR013785 Aldolase_TIM, |
| FV81RT002HT2FL | 2.8 | 2.0 | 3.1 | 1.8 | histidine ammonia-lyase-like | IPR001106 Aromatic_Lyase IPR008948 L-Aspartase-like |
| NPNHRC_12201 | 2.7 | 2.3 | 2.6 | 2.5 | (Hydra viridis specific) |  |

(1) Gene annotations by blast2go.

**Supplemental Table 2.** List of the genes differentially expressed between Hv_Sym and Hv_Apo and fold changes of expression level examined by microarray and qPCR.

| Probe name<br>(gene ID) | Microarray | qPCR | HS_Best Hit (1) | InterProScan (2) |
| --- | --- | --- | --- | --- |
|  | A99/Apo | A99/Apo |  |  |
| 5168 | 9.3 | 7.4 |  | IPR000157 TIR_dom<br>PTHR23097 Tumor necrosis factor<br>receptor superfamily member |
| 6508 | 6.7 | 2.9 |  | IPR011029:DEATH-like_dom |
| 11411 | 2.9 | 2.0 | C-type mannose receptor 2 | IPR000742 EG-like_dom<br>IPR001304 C-type_lectin |
| 26108 | 7.2 | 7.2 | ephrin type-A receptor 6 isoform a | no IPS match |
| rc_2417 | 5.4 | 3.5 |  | IPR000488 Death_domain |
| rc_24563 | 6.1 | 6.7 | Proline-rich transmembrane protein 1 | IPR007593 CD225/Dispanin_fam<br>PTHR14948 NG5 |
| rc_9398 | 6.2 | 5.4 | protein-kinase, interferon-inducible<br>double stranded RNA dependent<br>inhibitor, repressor of (P58 repressor) | PTHR11697 general transcription factor 2-related zinc finger protein |

A99<Apo
| Probe name<br>(gene ID) | Microarray | qPCR | HS_best hit | InterProScan |
| --- | --- | --- | --- | --- |
|  | Apo/A99 | Apo/A99 |  |  |
| rc_10789 | 2.5 | 3.7 | endoribonuclease Dicer | IPR000999 RNase_III_dom<br>PTHR1495 helicase-related |
| rc_12826 | 3.0 | 2.3 | interferon regulatory factor 1 | IPR001346 Interferon_reg_fact_DNA-bd_dom;<br>IPR011991 WHTH_DNA-bd_dom<br>PTHR11949 interferon regulatory factor |
| rc_8898 | 6.1 | 4.1 | leucine-rich repeat-containing protein 15<br>isoform b | IPR001611 Leu-rich_rp<br>PTHR24373 Toll-like receptor 9 |
| FV81RT001CSTY | 3.2 | 2.0 | astrocytic phosphoprotein PEA-15 | IPR001875 DED, IPR011029 DEATH-like_dom |
| RSASM_17752 | 4.0 | 2.1 | CD97 antigen isoform 2 precursor | IPR000832 GPCR_2_secretin-like<br>PTHR12011 vasoactive intestinal<br>polypeptide receptor 2 |

(1) Best hit human genes. (2) Results of domain search.

**Supplemental Table 3.** Summary of sequence data for assembling Chlorella sp. A99 genome sequences

|  |  |  |
| --- | --- | --- |
| **Number of reads** | 85469010 |  |
| **Number of reads assembled** | 61838513 |  |
| **Number of bases** | 17398635102 |  |
|  | **Scaffolds** | **Contigs** |
| **Total length of sequence** | 40934037 | 40687875 |
| **Total number of sequences** | 82 | 7455 |
| **Maximum length of sequence** | 4003385 | 171868 |
| **N50** | 1727419 | 12747 |
| **GC contents (%)** | 68.07 % | 69.95 % |

**Supplemental Table 4.** A. The number of Pfam domains related to amino acids transport B. Ortholog groups including Aa_trans containing genes C. Blast best hit genes of Arabidopsis thaliana of genes belonging to OG0000040 and OG0000324

A. The number of Pfam domains related to amino acids transport
| Pfam Domain Name | A99 | NC64A | c169 | Cr | Vc | Mp | Ot |
| --- | --- | --- | --- | --- | --- | --- | --- |
| Aa_trans | 30 | 38 | 21 | 9 | 7 | 9 | 8 |
| AA_permease | 4 | 6 | 15 | 5 | 6 | 1 | 1 |

B. Ortholog groups including Aa\_trans containing genes
| Ortholog Group ID | A99 | NC64A | c169 | Cr | Vc | Mp | Ot |
| --- | --- | --- | --- | --- | --- | --- | --- |
| OG0000040 | 12 | 12 | 6 | 3 | 1 | 0 | 0 |
| OG0000324 | 6 | 7 | 1 | 2 | 1 | 0 | 0 |
| OG0001336 | 2 | 1 | 1 | 1 | 1 | 1 | 1 |
| OG0004053 | 1 | 2 | 2 | 1 | 0 | 0 | 0 |
| OG0006517 | 1 | 1 | 0 | 0 | 0 | 0 | 0 |
| OG0001069 | 1 | 2 | 3 | 1 | 1 | 1 | 1 |
| OG0000830 | 1 | 4 | 1 | 2 | 1 | 2 | 1 |
| OG0002190 | 1 | 1 | 1 | 1 | 1 | 1 | 1 |
| OG0011340 | 1 | 0 | 0 | 0 | 0 | 0 | 0 |
| OG0004863 | 1 | 5 | 0 | 0 | 0 | 0 | 0 |
| OG0000468 | 2 | 2 | 2 | 2 | 2 | 2 | 2 |
| OG0003354 | 2 | 1 | 1 | 1 | 1 | 1 | 1 |
| OG0003801 | 1 | 1 | 0 | 1 | 1 | 1 | 1 |

C. Blast best hit genes of Arabidopsis thaliana of genes belonging to OG0000040 and OG0000324
| OG0000040 | Best Hit gene ID | Best Hit gene of Arabidopsis thaliana | e-value |
| --- | --- | --- | --- |
| scaffold1.g5447.t1 | NP_196484.1 | amino acid permease 2 | 4E-62 |
| scaffold1.g5579.t1 | NP_196484.1 | amino acid permease 2 | 7E-32 |
| scaffold12.g8277.t1 | NP_196484.1 | amino acid permease 2 | 1E-34 |
| scaffold13.g380.t1 | NP_196484.1 | amino acid permease 2 | 6E-12 |
| scaffold14.g1284.t1 | NP_196484.1 | amino acid permease 2 | 2E-33 |
| scaffold2.g7119.t1 | NP_001318716.1 | lysine histidine transporter 1 | 9E-73 |
| scaffold2.g7251.t1 | NP_196484.1 | amino acid permease 2 | 3E-59 |
| scaffold21.g2221.t1 | NP_175076.2 | amino acid permease 5 | 9E-60 |
| scaffold40.g5168.t1 | NP_196484.1 | amino acid permease 2 | 5E-38 |
| scaffold5.g864.t1 | NP_196484.1 | amino acid permease 2 | 2E-60 |
| scaffold6.g2644.t1 | NP_196484.1 | amino acid permease 2 | 2E-58 |
| scaffold6.g2815.t1 | NP_186825.2 | Transmembrane amino acid transporter family protein | 3E-43 |

| **OG0000324** | **Best Hit gene ID** | **Best Hit gene of *Arabidopsis thaliana*** | **e-value** |
| --- | --- | --- | --- |
| scaffold10.g2481.t1 | NP_001330273.1 | Transmembrane amino acid transporter family protein | 7E-06 |
| scaffold11.g3916.t1* | NP_172258.1 | Histone superfamily protein | 1E-47 |
|  | NP_565239.1 | Transmembrane amino acid transporter family protein | 7E-40 |
| scaffold15.g4364.t1 | NP_566854.1 | Transmembrane amino acid transporter family protein | 8E-31 |
| scaffold2.g7197.t1 | NP_566854.1 | Transmembrane amino acid transporter family protein | 2E-24 |
| scaffold21.g2185.t1 | NP_565239.1 | Transmembrane amino acid transporter family protein | 5E-14 |
| scaffold3.g6325.t1 | NP_566854.1 | Transmembrane amino acid transporter family protein | 6E-34 |

\* This sequence consists from a region similar to NP\_172258.1 and a region similar to NP\_565239.1.

**Supplementary Table 5.**
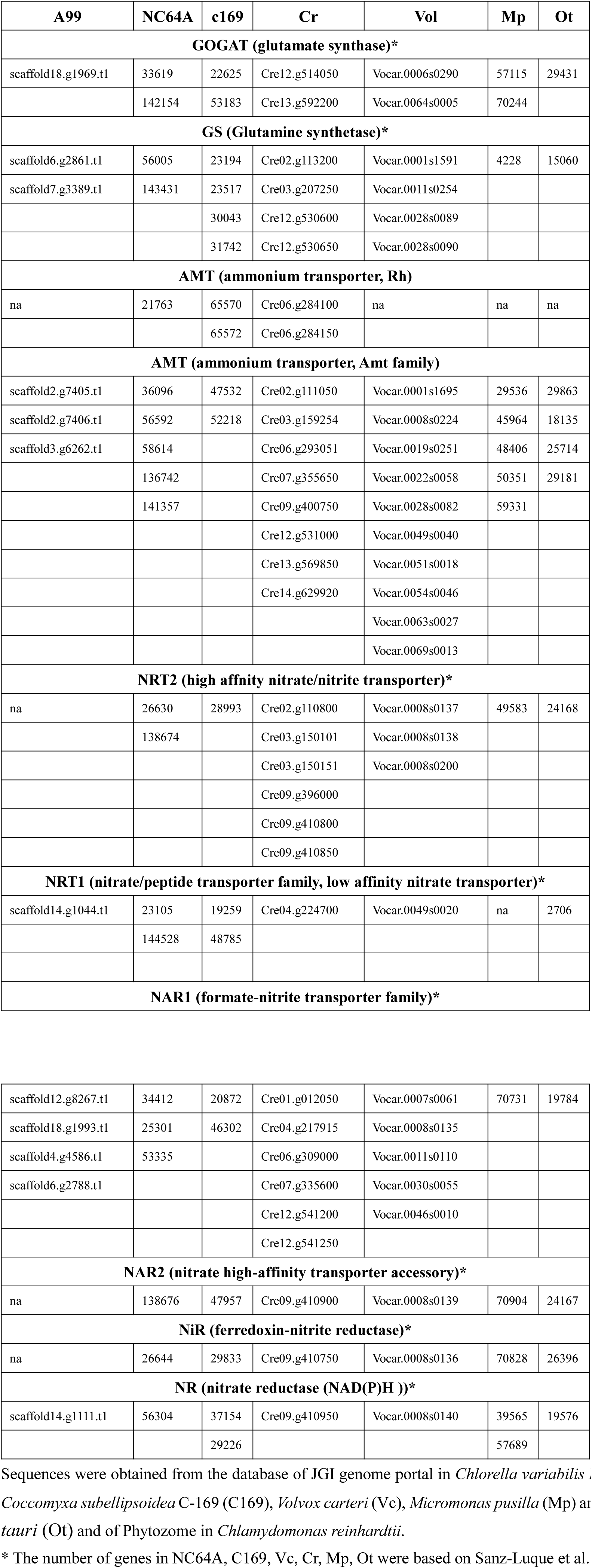
Sequence ID of nitrogen assimilation genes

**Supplemental Table 6.**
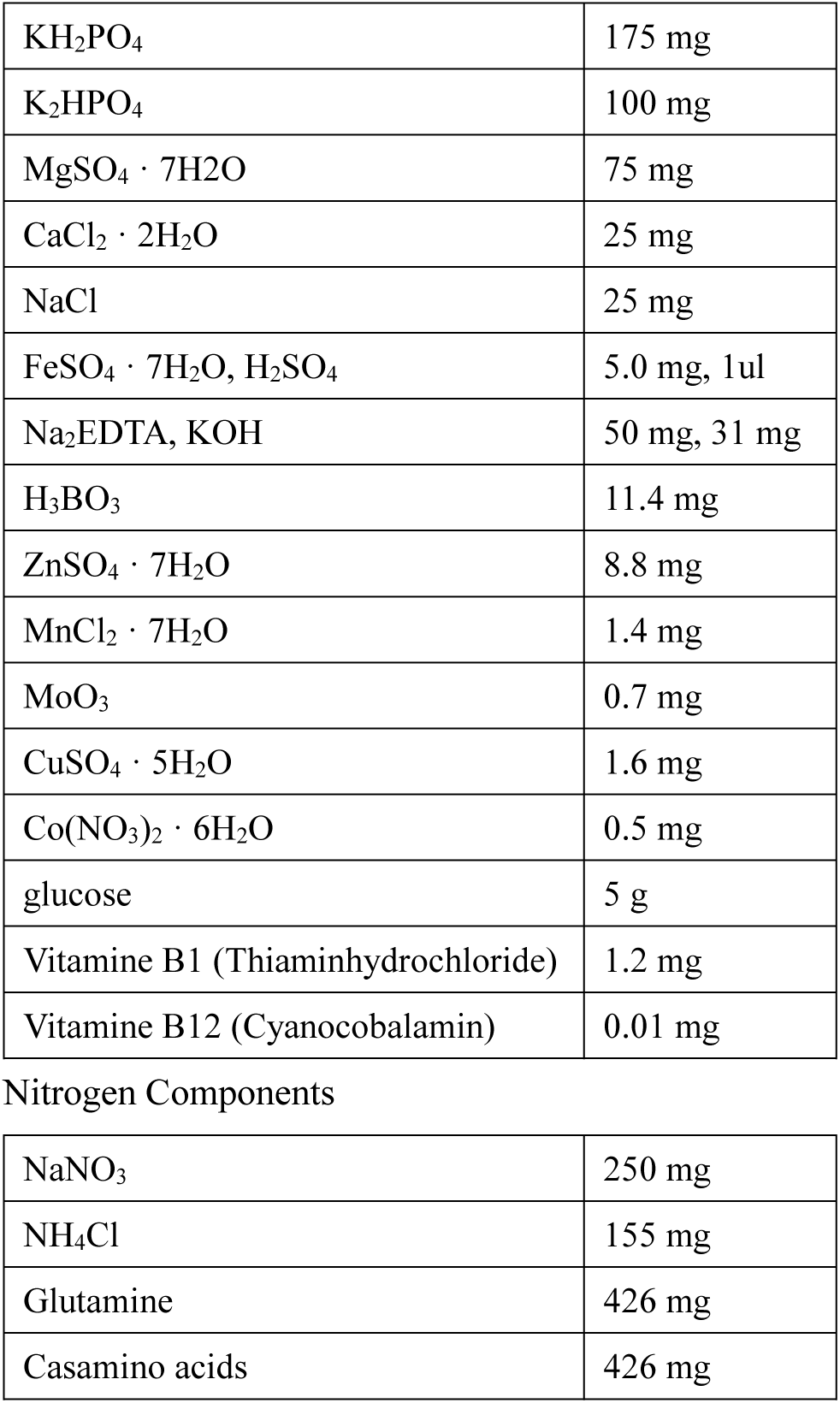
Composition of modified Bold’s Basal Medium for 1 liter (pH. 7)

|  |  |
| --- | --- |
| $\text{KH}_2\text{PO}_4$ | 175 mg |
| $\text{K}_2\text{HPO}_4$ | 100 mg |
| $\text{MgSO}_4 \cdot 7\text{H}_2\text{O}$ | 75 mg |
| $\text{CaCl}_2 \cdot 2\text{H}_2\text{O}$ | 25 mg |
| $\text{NaCl}$ | 25 mg |
| $\text{FeSO}_4 \cdot 7\text{H}_2\text{O}$, $\text{H}_2\text{SO}_4$ | 5.0 mg, 1ul |
| $\text{Na}_2\text{EDTA}$, $\text{KOH}$ | 50 mg, 31 mg |
| $\text{H}_3\text{BO}_3$ | 11.4 mg |
| $\text{ZnSO}_4 \cdot 7\text{H}_2\text{O}$ | 8.8 mg |
| $\text{MnCl}_2 \cdot 7\text{H}_2\text{O}$ | 1.4 mg |
| $\text{MoO}_3$ | 0.7 mg |
| $\text{CuSO}_4 \cdot 5\text{H}_2\text{O}$ | 1.6 mg |
| $\text{Co}(\text{NO}_3)_2 \cdot 6\text{H}_2\text{O}$ | 0.5 mg |
| glucose | 5 g |
| Vitamine B1 (Thiaminhydrochloride) | 1.2 mg |
| Vitamine B12 (Cyanocobalamin) | 0.01 mg |
| Nitrogen Components |  |
| $\text{NaNO}_3$ | 250 mg |
| $\text{NH}_4\text{Cl}$ | 155 mg |
| Glutamine | 426 mg |
| Casamino acids | 426 mg |

**Supplemental Table 7.**
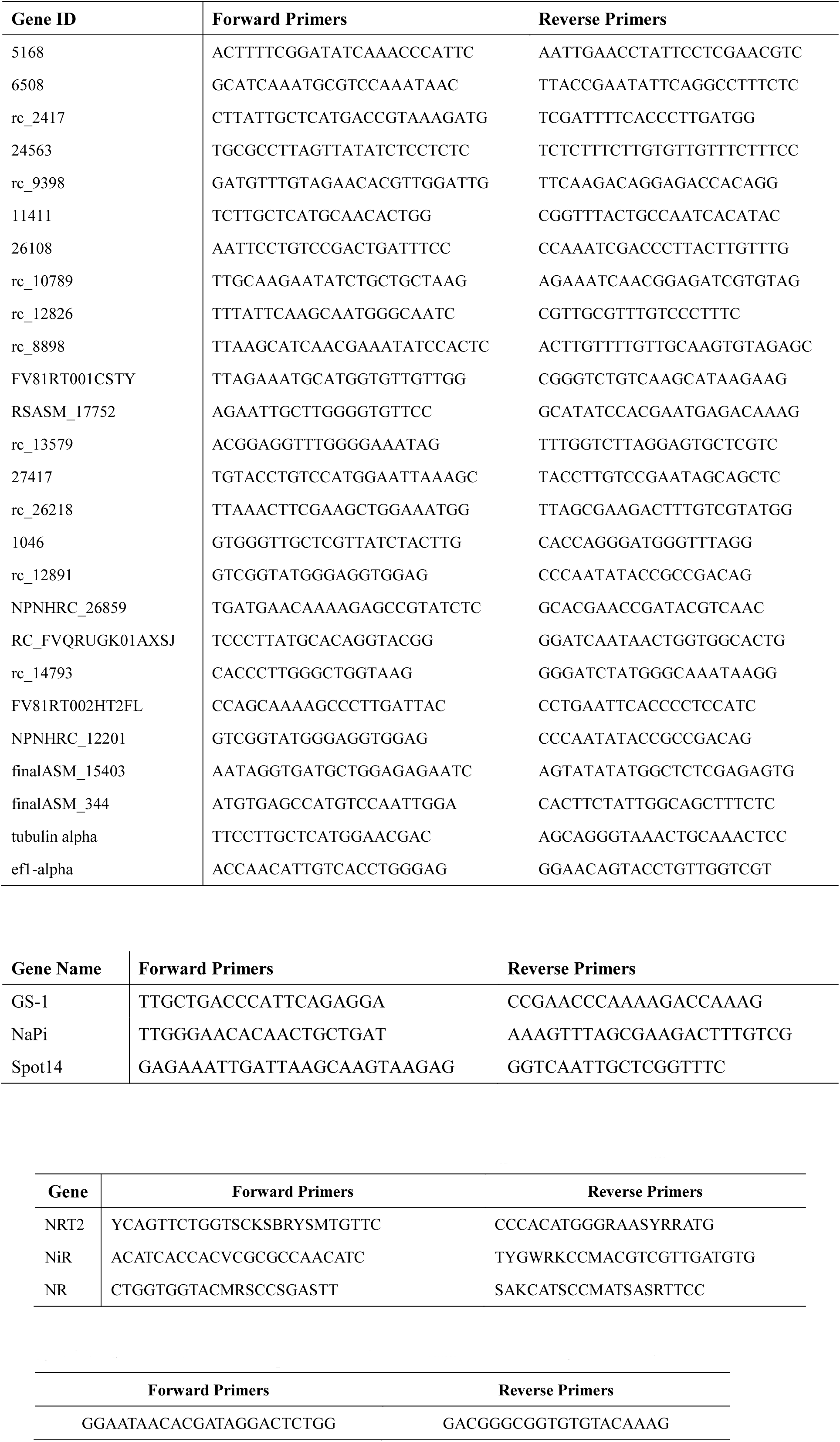
A. Primer sequences for quantitative real time RT-PCR B. Primer sequences for *in situ* hybridization probes C. Primer sequences for PCR amplification of nitrogen assimilation genes in green algae D. Primer sequences for PCR amplification of 18S ribosomal DNA gene in green algae

## Supplementary Figures

**Supplementary Figure 1.**
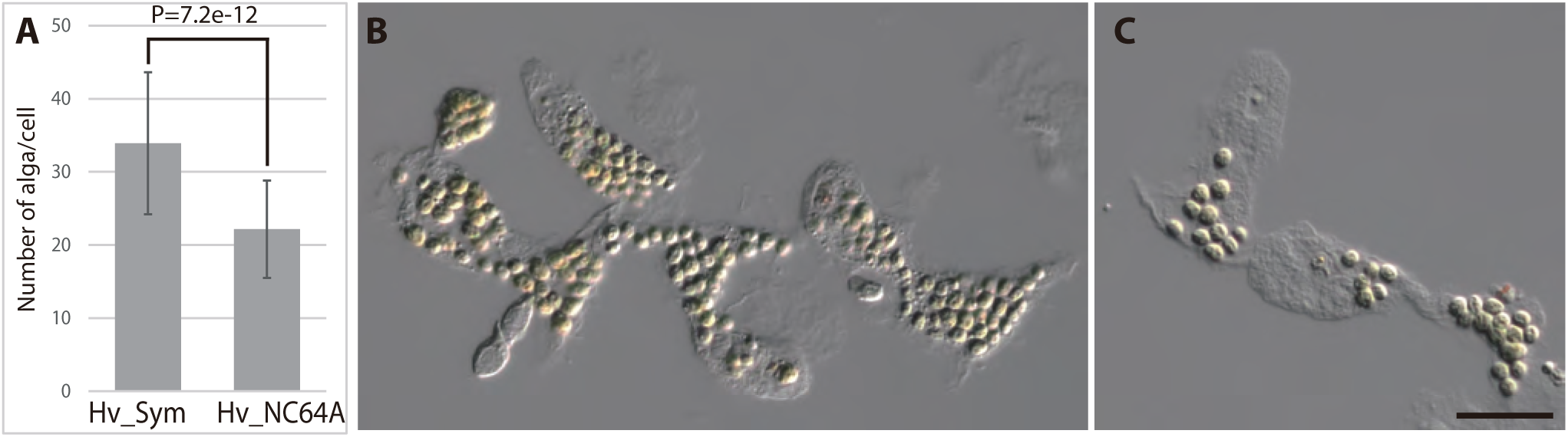
*Chlorella* sp. A99 and *Chlorella variabilis* NC64A in *Hydra viridissima* A99. (A) Average number of algae per *Hydra* cell, for native *Chlorella* sp. A99 (Hv_Sym) and aposymbiont *Hydra* re-infected with *Chlorella variabilis* NC64A (Hv_NC64A). (B) Endodermal epithelial cells of Hv_Sym showing intracellular algae (C) Endodermal epithelial cells of Hv_NC64A. Scale bar, 20 μm

**Supplementary Figure 2.**
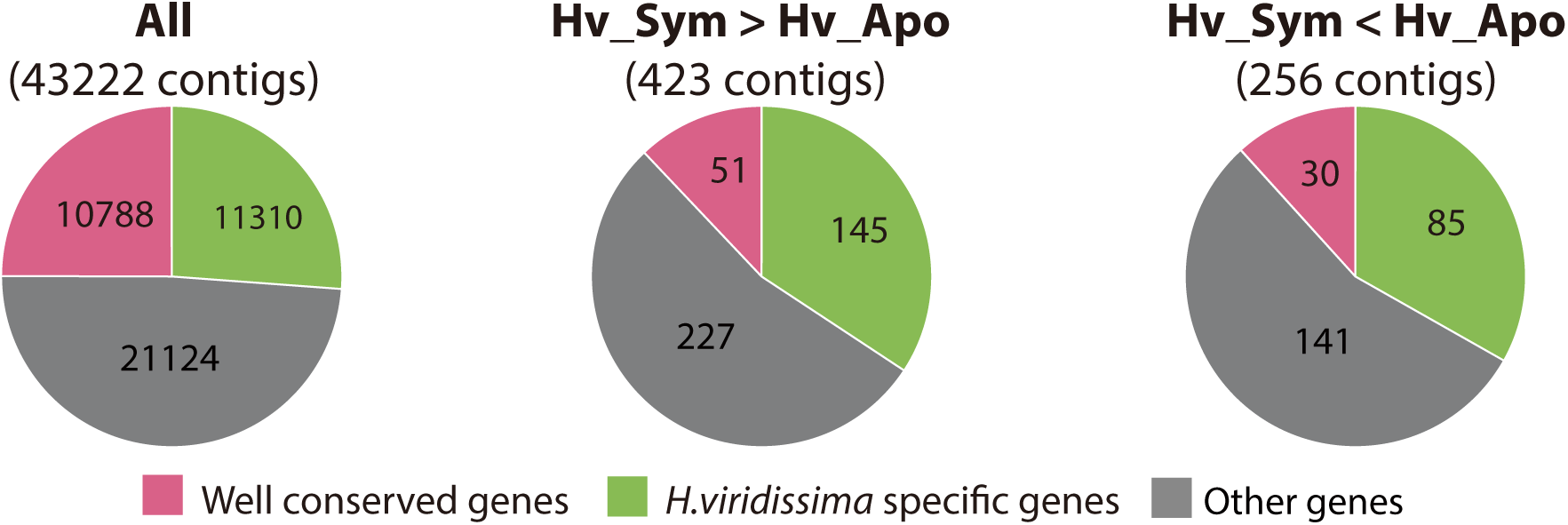
Conserved genes and species-specific genes differentially expressed in symbiotic *Hydra*. Distribution of well-conserved *Hydra viridissma* genes (pink), *Hydra viridissima*-specific genes (green) and other genes (shared by some but not all metazoans, gray) among eight metazoans: *Hydra magnipapillata, Acropora digitifera, Nematostella vectensis, Strongylocentrotus pupuratus, Branchiostoma floridae, Homo sapiens* and *Drosophila melanogaster* and *Hydra viridissima*. Pie charts are shown for all contigs (All), up-regulated contigs (Hv_Sym > Hv_Apo) and down-regulated contigs (Hv_Sym < Hv_Apo).

**Supplementary Figure 3.**
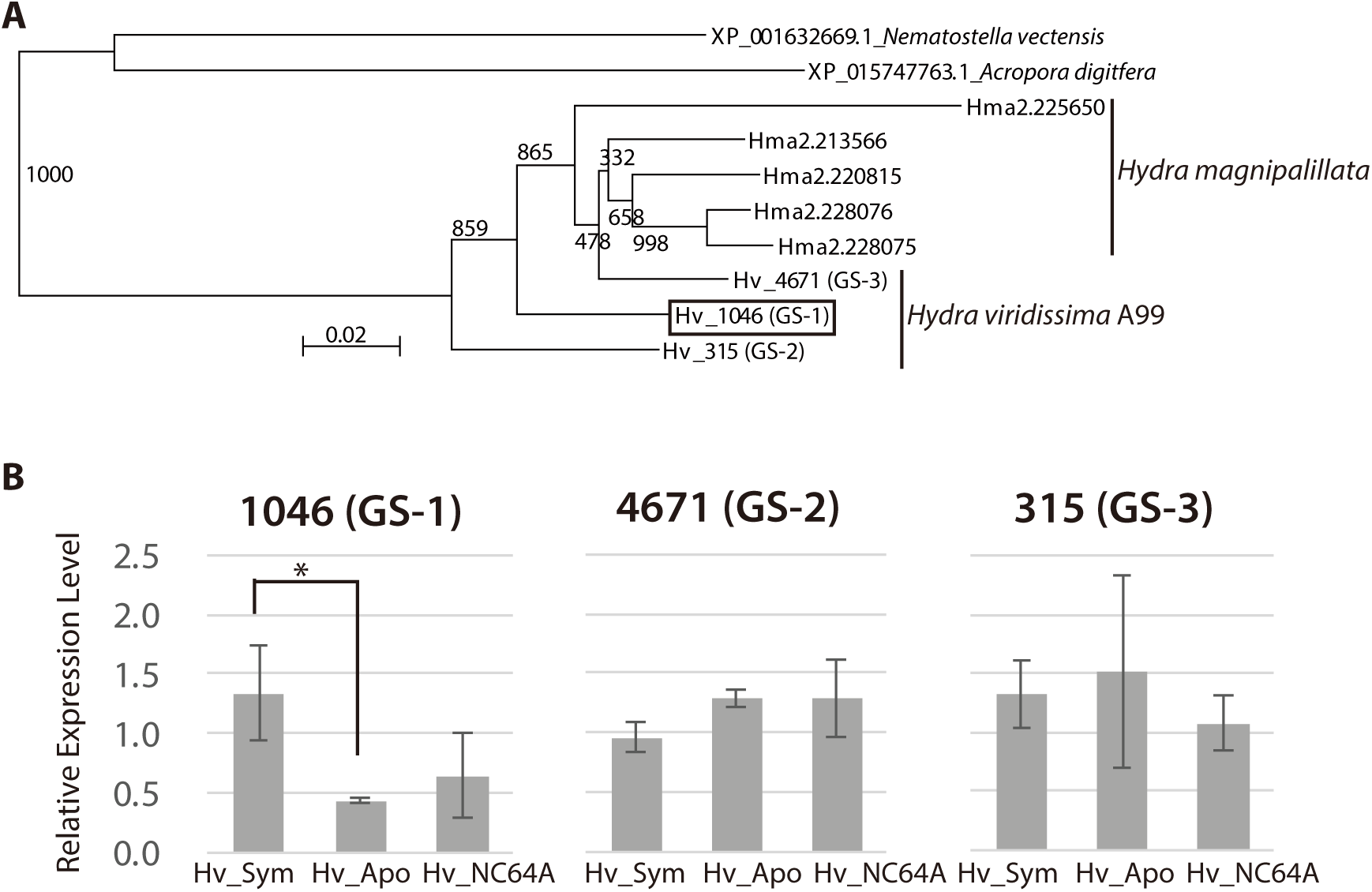
Glutamine synthetase (GS) genes in Cnidarians. (A) Phylogenetic tree of the GS gene of four species in Cnidarians. While anthozoans (*Nematostella vectensis, Acropora digitifera*) have a single GS gene, *Hydra magnipappilata* (Hma) has five genes and *Hydra viridissima* A99 has three genes, Hv_1046 (GS-1), Hv_315 (GS-2) and Hv_4671 (GS-3). (B) Relative expression level of the three GS genes in Hv_Sym, Hv_NC64A and Hv_Apo as determined by microarray analysis.

**Supplementary Figure 4.**
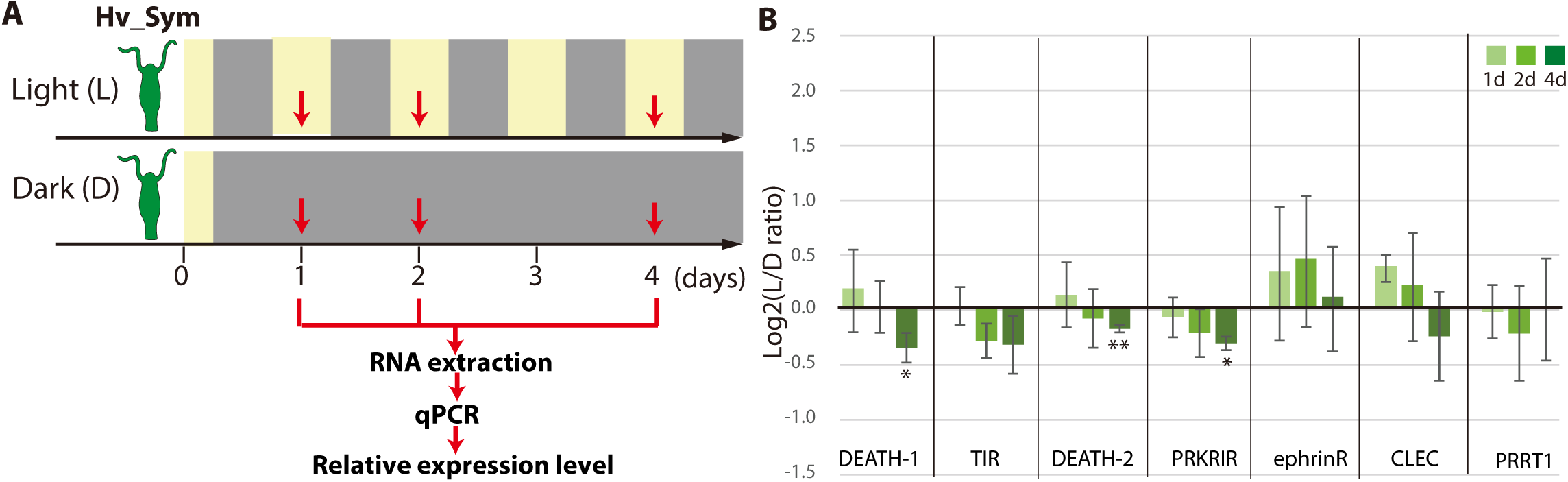
Differential expression of symbiosis-dependent *Hydra* genes grown under light/dark condition and in darkness. (A) Sampling scheme. Hv_Sym was cultured in the light-dark condition (Light: L) and in the continuous dark (Dark: D). Gene expression levels were examined by qPCR at 1, 2, 4 days for each condition (red arrows). (B) Expression difference of the genes in Hv_A99 between the two conditions. DEATH-1 and DEATH-2: Death domain containing proteins (gene ID: 6508 and rc_2417), TIR: Toll/interleukin-1 receptor domain containing protein (gene ID: 5168), PRKRIR: protein-kinase interferon-inducible double stranded RNA dependent inhibitor, repressor of (p58 repressor) (gene ID: rc_9398), ephrinR: ephrin receptor (gene ID: 26108), CLEC: C-type mannose receptor (gene ID: 11411), PRRT1: proline-rich transmembrane protein 1 (gene ID: rc_24563). The vertical axis shows log scale (log2) fold change of relative expression levels in the light condition over the dark condition. T-test evalue, * <0.05, ** <0.01.

**Supplementary Figure 5.**
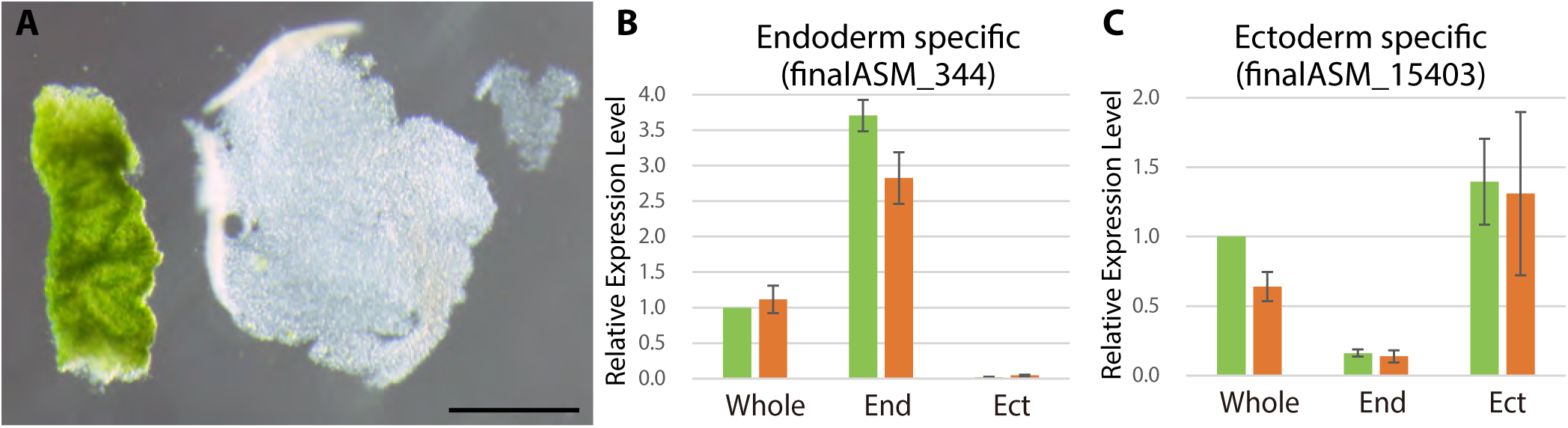
Tissue isolation of green hydra. (A) Isolated endoderm (left) and isolated ectoderm (right). Scale bar, 1 mm. Expression levels of an endoderm specific gene finalASM_15403 (B) and that of an ectoderm specific gene finalASM_344 (C) in whole hydra (Whole) and isolated endoderm (End) and ectoderm (Ect) were examined to confirm whether tissue isolation had performed properly.

**Supplementary Figure 6.**
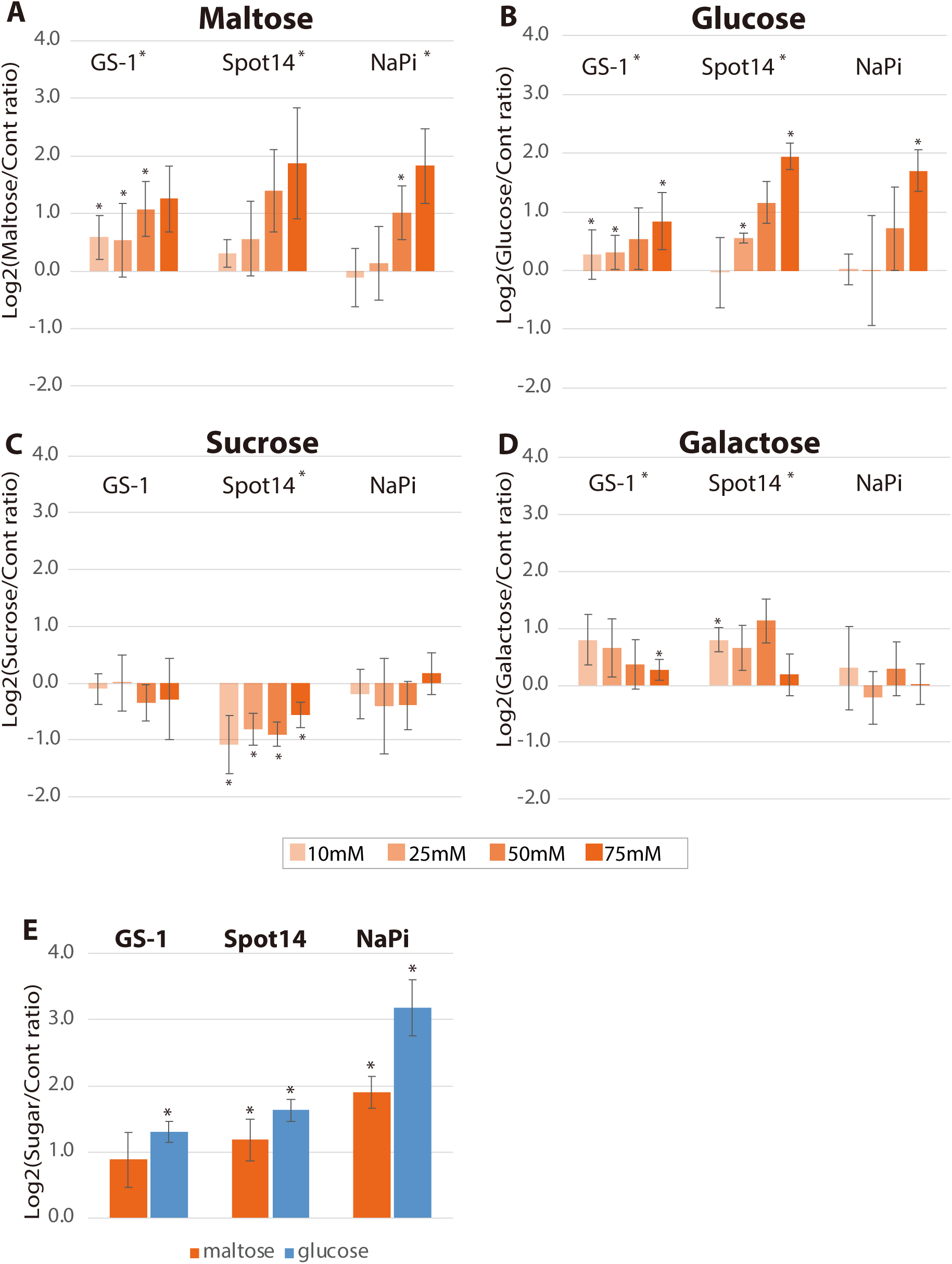
Effects of sugars on *Hydra* growth. Effects of growth in presence of maltose (A), glucose (B), sucrose (C) and galactose (D) on gene expression of GS-1, Spot14 and NaPi. Hv_Apo were cultured in medium containing 10 mM, 25 mM, 50 mM or 75 mM of each sugar for 48 hours, and 75 mM maltose (orange) and glucose (blue) for 6 hours (E). RNA was extracted from the polyps in the light condition. Expression difference of the genes was examined by qPCR. The vertical axis is log scale (log2) fold change of relative expression level of sugar-treated hydras over controls. Error bars indicate standard deviation. T-test in each concentration and Kruskal-Wallis test in the series of 48 hours treatment were performed. * p-value <0.05

**Supplementary Figure 7.**
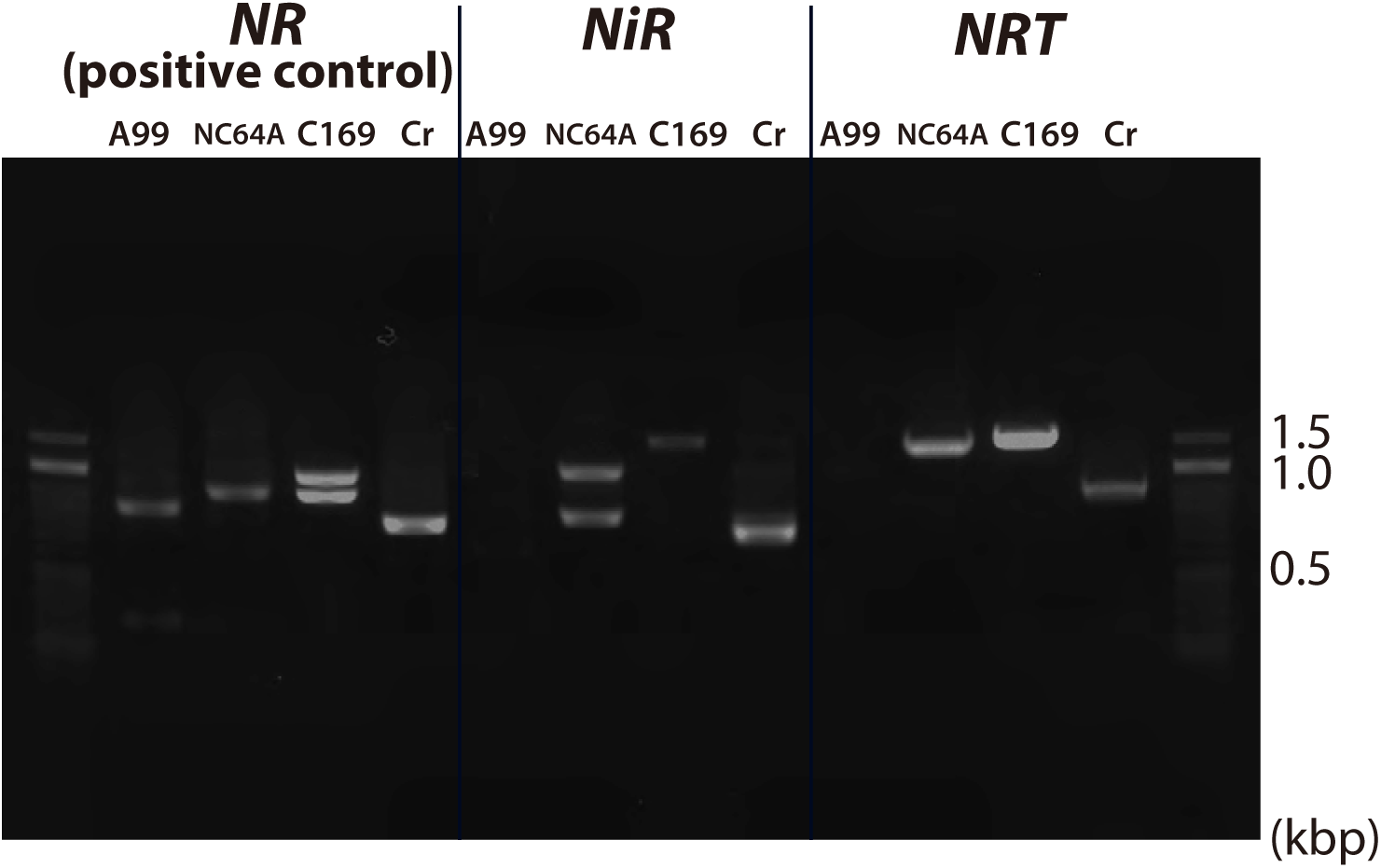
PCR of nitrate assimilation genes. PCR amplification of genomic DNA corresponding to the genes *NRT2, NiR* and *NR* (positive control) was performed in *Chlorella* sp. A99 (A99), *Chlorella variabilis* NC64A (NC64A), *Coccomyxa subellipsoidea* C169 (C169) and *Chlamydomonas reinhardtii* (Cr).

**Supplementary Figure 8.**
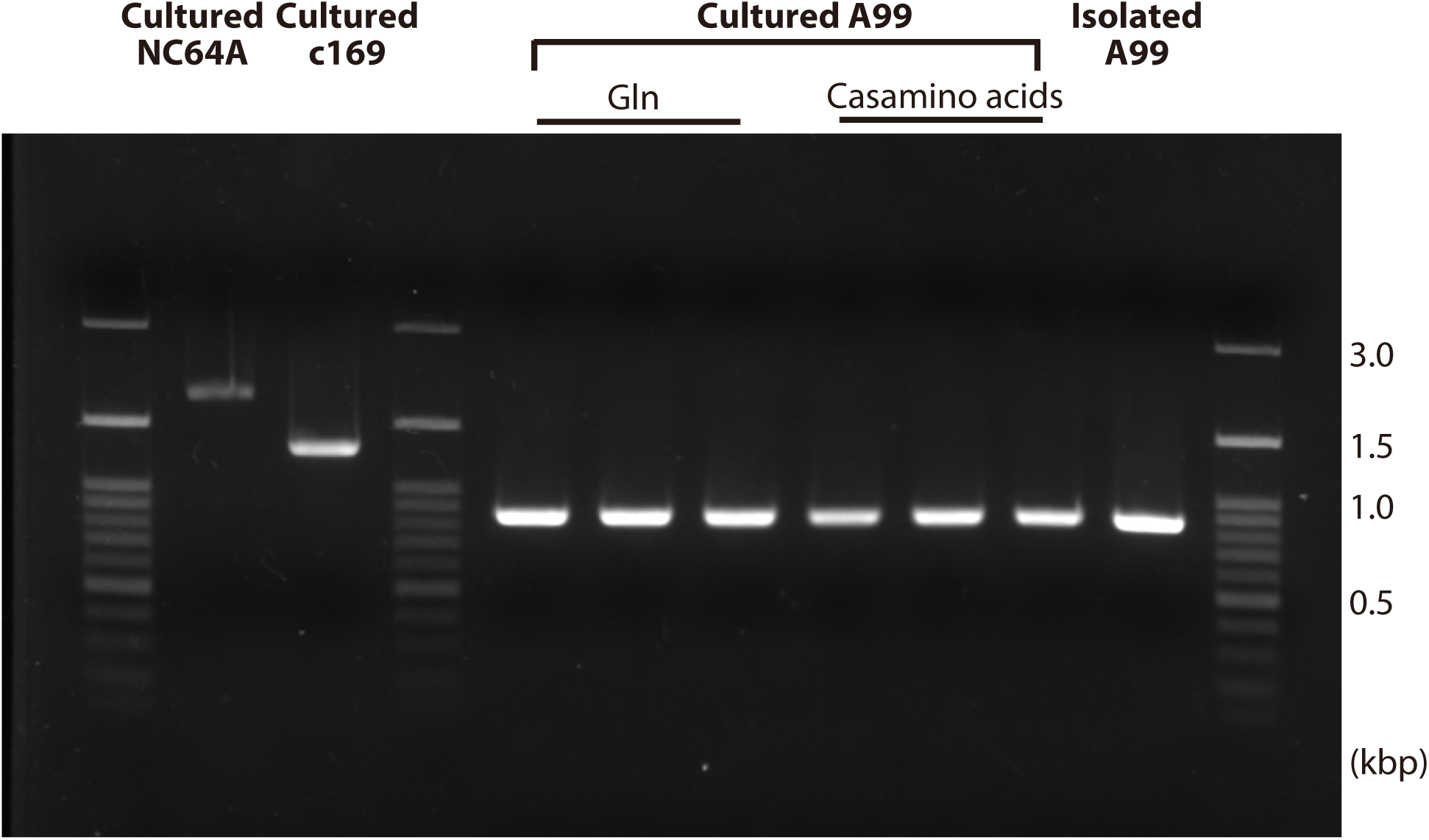
PCR of 18S rRNA genes in cultured algae. PCR amplification of genomic DNA of the 18S rRNA gene was performed in *Chlorella* A99 shortly after isolation from *H. viridissima* A99 (Isolated A99), cultured in medium containing glutamine (Glu) and in medium with casamino acids for 12 days, with cultured NC64A and C169 added for comparison.

## References

Baumgarten, S., Simakov, O., Esherick, L.Y., Liew, Y.J., Lehnert, E.M., Michell, C.T., Li, Y., Hambleton, E.A., Guse, A., Oates, M.E., Gough, J., Weis, V.M., Aranda, M., Pringle, J.R., Voolstra, C.R., 2015. The genome of Aiptasia, a sea anemone model for coral symbiosis. Proceedings of the National Academy of Sciences 112, 11893–11898. 10.1073/pnas.1513318112

Bischoff, H.W., Bold, H.C., 1963. Some soil algae from enchanted rock and related Algal species. [s.n.], Austin.

Blanc, G., Agarkova, I., Grimwood, J., Kuo, A., Brueggeman, A., Dunigan, D.D., Gurnon, J., Ladunga, I., Lindquist, E., Lucas, S., Pangilinan, J., Proschold, T., Salamov, A., Schmutz, J., Weeks, D., Yamada, T., Lomsadze, A., Borodovsky, M., Claverie, J.M., Grigoriev, I.V., Van Etten, J.L., 2012. The genome of the polar eukaryotic microalga Coccomyxa subellipsoidea reveals traits of cold adaptation. Genome Biol. 13, R39. 10.1186/gb-2012-13-5-r39

Blanc, G., Duncan, G., Agarkova, I., Borodovsky, M., Gurnon, J., Kuo, A., Lindquist, E., Lucas, S., Pangilinan, J., Polle, J., Salamov, A., Terry, A., Yamada, T., Dunigan, D.D., Grigoriev, I.V., Claverie, J.M., Van Etten, J.L., 2010. The Chlorella variabilis NC64A genome reveals adaptation to photosymbiosis, coevolution with viruses, and cryptic sex. Plant Cell 22, 2943–2955. 10.1105/tpc.110.076406

Boetzer, M., Henkel, C.V., Jansen, H.J., Butler, D., Pirovano, W., 2011. Scaffolding pre-assembled contigs using SSPACE. Bioinformatics 27, 578–579. 10.1093/bioinformatics/btq683

Bosch, T.C., Augustin, R., Anton-Erxleben, F., Fraune, S., Hemmrich, G., Zill, H., Rosenstiel, P., Jacobs, G., Schreiber, S., Leippe, M., Stanisak, M., Grotzinger, J., Jung, S., Podschun, R., Bartels, J., Harder, J., Schroder, J.M., 2009. Uncovering the evolutionary history of innate immunity: the simple metazoan Hydra uses epithelial cells for host defence. Dev. Comp. Immunol. 33, 559–569. 10.1016/j.dci.2008.10.004

Bossert, P., Dunn, K.W., 1986. Regulation of intracellular algae by various strains of the symbiotic Hydra viridissima. J. Cell Sci. 85, 187–195

Brown, S.B., Maloney, M., Kinlaw, W.B., 1997. “Spot 14” protein functions at the pretranslational level in the regulation of hepatic metabolism by thyroid hormone and glucose. J. Biol. Chem. 272, 2163–2166

Campbell, R.D., 1990. Transmission of symbiotic algae through sexual reproduction in hydra: movement of algae into the oocyte. Tissue Cell 22, 137–147

Cernichiari, E., Muscatine, L., Smith, D.C., 1969. Maltose Excretion by the Symbiotic Algae of Hydra viridis. Proc. R. Soc. Lond., Ser. B: Biol. Sci. 173, 557–576. 10.1098/rspb.1969.0077

Conesa, A., Gotz, S., Garcia-Gomez, J.M., Terol, J., Talon, M., Robles, M., 2005. Blast2GO: a universal tool for annotation, visualization and analysis in functional genomics research. Bioinformatics 21, 3674–3676. 10.1093/bioinformatics/bti610

Cook, C.B., Kelty, M.O., 1982. Glycogen, protein, and lipid content of green, aposymbiotic, and nonsymbiotic hydra during starvation. J. Exp. Zool. 222, 1–9. 10.1002/jez.1402220102

Davy, S.K., Allemand, D., Weis, V.M., 2012a. Cell Biology of Cnidarian-Dinoflagellate Symbiosis. Microbiology and Molecular Biology Reviews: MMBR 76, 229–261. 10.1128/MMBR.05014-11

Davy, S.K., Allemand, D., Weis, V.M., 2012b. Cell biology of cnidarian-dinoflagellate symbiosis. Microbiol. Mol. Biol. Rev. 76, 229–261. 10.1128/MMBR.05014-11

Dorling, M., McAuley, P.J., Hodge, H., 1997. Effect of pH on growth and carbon metabolism of maltose-releasingChlorella(Chlorophyta). Eur. J. Phycol. 32, 19–24. 10.1080/09541449710001719335

Douglas, A., Smith, D.C., 1984. The Green Hydra Symbiosis. VIII. Mechanisms in Symbiont Regulation. Proceedings of the Royal Society of London. Series B. Biological Sciences 221, 291–319. 10.1098/rspb.1984.0035

Douglas, A.E., 1994. Symbiotic interactions/by A.E. Douglas. Oxford University Press, Oxford; New York.

Douglas, A.E., Smith, D.C., 1983. The cost of symbionts to the host in the green hydra symbiosis. W. DeGriiyter and Co. Berlin.

Eddy, S.R., 1998. Profile hidden Markov models. Bioinformatics 14, 755–763

Emms, D.M., Kelly, S., 2015. OrthoFinder: solving fundamental biases in whole genome comparisons dramatically improves orthogroup inference accuracy. Genome Biol. 16, 157. 10.1186/s13059-015-0721-2

Finn, R.D., Coggill, P., Eberhardt, R.Y., Eddy, S.R., Mistry, J., Mitchell, A.L., Potter, S.C., Punta, M., Qureshi, M., Sangrador-Vegas, A., Salazar, G.A., Tate, J., Bateman, A., 2016. The Pfam protein families database: towards a more sustainable future. Nucleic Acids Res. 44, D279–285. 10.1093/nar/gkv1344

Gao, C., Wang, Y., Shen, Y., Yan, D., He, X., Dai, J., Wu, Q., 2014. Oil accumulation mechanisms of the oleaginous microalga Chlorella protothecoides revealed through its genome, transcriptomes, and proteomes. BMC Genomics 15, 582. 10.1186/1471-2164-15-582

Habetha, M., Anton-Erxleben, F., Neumann, K., Bosch, T.C., 2003. The Hydra viridis/Chlorella symbiosis. Growth and sexual differentiation in polyps without symbionts. Zoology 106, 101–108. 10.1078/0944-2006-00104

Habetha, M., Bosch, T.C., 2005. Symbiotic Hydra express a plant-like peroxidase gene during oogenesis. J. Exp. Biol. 208, 2157–2165. 10.1242/jeb.01571

Hall, 1999. BioEdit: a user-friendly biological sequence alignment editor and analysis program for Windows 95/98/NT. Nucleic Acid Symposium Series 41, 95–98

Huss, V.A.R., Holweg, C., Seidel, B., Reich, V., Rahat, M., Kessler, E., 1993/1994. There is an ecological basis for host/symbiont specificity in Chlorella/Hydra symbioses. Endocytobiosis and Cell Research 10, 35–46

Ishikawa, M., Yuyama, I., Shimizu, H., Nozawa, M., Ikeo, K., Gojobori, T., 2016. Different Endosymbiotic Interactions in Two Hydra Species Reflect the Evolutionary History of Endosymbiosis. Genome Biol. Evol. 8, 2155–2163. 10.1093/gbe/evw142

Joy, J.B., 2013. Symbiosis catalyses niche expansion and diversification. Proceedings of the Royal Society B: Biological Sciences 280. 10.1098/rspb.2012.2820

Kamako, S.-i., Hoshina, R., Ueno, S., Imamura, N., 2005. Establishment of axenic endosymbiotic strains of Japanese Paramecium bursaria and the utilization of carbohydrate and nitrogen compounds by the isolated algae. Eur. J. Protistol. 41, 193–202. 10.1016/j.ejop.2005.04.001

Kanehisa, M., Goto, S., 2000. KEGG: kyoto encyclopedia of genes and genomes. Nucleic Acids Res. 28, 27–30

Karakashian, S.J., Karakashian, M.W., 1965. Evolution and Symbiosis in the Genus Chlorella and Related Algae. Evolution 19, 368–377

Kawaida, H., Ohba, K., Koutake, Y., Shimizu, H., Tachida, H., Kobayakawa, Y., 2013. Symbiosis between hydra and chlorella: molecular phylogenetic analysis and experimental study provide insight into its origin and evolution. Mol. Phylogenet. Evol. 66, 906–914. 10.1016/j.ympev.2012.11.018

Khalturin, K., Hemmrich, G., Fraune, S., Augustin, R., Bosch, T.C., 2009. More than just orphans: are taxonomically-restricted genes important in evolution? Trends Genet. 25, 404–413. 10.1016/j.tig.2009.07.006

Kishimoto, Y., Murate, M., Sugiyama, T., 1996. Hydra regeneration from recombined ectodermal and endodermal tissue. I. Epibolic ectodermal spreading is driven by cell intercalation. J. Cell Sci. 109, 763–772

Krapp, A., David, L.C., Chardin, C., Girin, T., Marmagne, A., Leprince, A.-S., Chaillou, S., Ferrario-Méry, S., Meyer, C., Daniel-Vedele, F., 2014. Nitrate transport and signalling in Arabidopsis. J. Exp. Bot. 65, 789–798. 10.1093/jxb/eru001

Larkin, M.A., Blackshields, G., Brown, N.P., Chenna, R., McGettigan, P.A., McWilliam, H., Valentin, F., Wallace, I.M., Wilm, A., Lopez, R., Thompson, J.D., Gibson, T.J., Higgins, D.G., 2007. Clustal W and Clustal X version 2.0. Bioinformatics 23, 2947–2948. 10.1093/bioinformatics/btm404

Lehnert, E.M., Mouchka, M.E., Burriesci, M.S., Gallo, N.D., Schwarz, J.A., Pringle, J.R., 2014. Extensive Differences in Gene Expression Between Symbiotic and Aposymbiotic Cnidarians. G3: Genes|Genomes|Genetics 4, 277–295. 10.1534/g3.113.009084

Liaw, S.-H., Kuo, I., Eisenberg, D., 1995. Discovery of the ammonium substrate site on glutamine synthetase, A third cation binding site. Protein Sci. 4, 2358–2365. 10.1002/pro.5560041114

Luo, R., Liu, B., Xie, Y., Li, Z., Huang, W., Yuan, J., He, G., Chen, Y., Pan, Q., Liu, Y., Tang, J., Wu, G., Zhang, H., Shi, Y., Liu, Y., Yu, C., Wang, B., Lu, Y., Han, C., Cheung, D.W., Yiu, S.M., Peng, S., Xiaoqian, Z., Liu, G., Liao, X., Li, Y., Yang, H., Wang, J., Lam, T.W., Wang, J., 2012. SOAPdenovo2: an empirically improved memory-efficient short-read de novo assembler. Gigascience 1, 18. 10.1186/2047-217X-1-18

Marcais, G., Kingsford, C., 2011. A fast, lock-free approach for efficient parallel counting of occurrences of k-mers. Bioinformatics 27, 764–770. 10.1093/bioinformatics/btr011

Martinez, D.E., Iniguez, A.R., Percell, K.M., Willner, J.B., Signorovitch, J., Campbell, R.D., 2010. Phylogeny and biogeography of Hydra (Cnidaria: Hydridae) using mitochondrial and nuclear DNA sequences. Mol. Phylogenet. Evol. 57, 403–410. 10.1016/j.ympev.2010.06.016

McAuley, P.J., 1986a. The cell cycle of symbiotic Chlorella. III. Numbers of algae in green hydra digestive cells are regulated at digestive cell division. J. Cell Sci. 85, 63–71

McAuley, P.J., 1986b. Isolation of viable uncontaminated Chlorella from green hydra. Limnol. Oceanogr. 31, 222–224. 10.4319/lo.1986.31.1.0222

McAuley, P.J., 1987a. Nitrogen limitation and amino-acid metabolism of Chlorella symbiotic with green hydra. Planta 171, 532–538. 10.1007/bf00392303

McAuley, P.J., 1987b. Quantitative Estimation of Movement of an Amino Acid from Host to Chlorella Symbionts in Green Hydra. Biol. Bull. 173, 504–512. 10.2307/1541696

McAuley, P.J., 1991. Amino acids as a nitrogen source for Chlorella symbiotic with green hydra, in: Williams, R.B., Cornelius, P.F.S., Hughes, R.G., Robson, E.A. (Eds.), Coelenterate Biology: Recent Research on Cnidaria and Ctenophora: Proceedings of the Fifth International Conference on Coelenterate Biology, 1989. Springer Netherlands, Dordrecht, pp. 369–376. 10.1007/978-94-011-3240-4_53

McAuley, P.J., Smith, D.C., 1982. The Green Hydra Symbiosis. V. Stages in the Intracellular Recognition of Algal Symbionts by Digestive Cells. Proc. R. Soc. Lond., Ser. B: Biol. Sci. 216, 7–23

McFall-Ngai, M., Hadfield, M.G., Bosch, T.C.G., Carey, H.V., Domazet-Lošo, T., Douglas, A.E., Dubilier, N., Eberl, G., Fukami, T., Gilbert, S.F., Hentschel, U., King, N., Kjelleberg, S., Knoll, A.H., Kremer, N., Mazmanian, S.K., Metcalf, J.L., Nealson, K., Pierce, N.E., Rawls, J.F., Reid, A., Ruby, E.G., Rumpho, M., Sanders, J.G., Tautz, D., Wernegreen, J.J., 2013. Animals in a bacterial world, a new imperative for the life sciences. Proceedings of the National Academy of Sciences 110, 3229–3236. 10.1073/pnas.1218525110

Merchant, S.S., Prochnik, S.E., Vallon, O., Harris, E.H., Karpowicz, S.J., Witman, G.B., Terry, A., Salamov, A., Fritz-Laylin, L.K., Marechal-Drouard, L., Marshall, W.F., Qu, L.H., Nelson, D.R., Sanderfoot, A.A., Spalding, M.H., Kapitonov, V.V., Ren, Q., Ferris, P., Lindquist, E., Shapiro, H., Lucas, S.M., Grimwood, J., Schmutz, J., Cardol, P., Cerutti, H., Chanfreau, G., Chen, C.L., Cognat, V., Croft, M.T., Dent, R., Dutcher, S., Fernandez, E., Fukuzawa, H., Gonzalez-Ballester, D., Gonzalez-Halphen, D., Hallmann, A., Hanikenne, M., Hippler, M., Inwood, W., Jabbari, K., Kalanon, M., Kuras, R., Lefebvre, P.A., Lemaire, S.D., Lobanov, A.V., Lohr, M., Manuell, A., Meier, I., Mets, L., Mittag, M., Mittelmeier, T., Moroney, J.V., Moseley, J., Napoli, C., Nedelcu, A.M., Niyogi, K., Novoselov, S.V., Paulsen, I.T., Pazour, G., Purton, S., Ral, J.P., Riano-Pachon, D.M., Riekhof, W., Rymarquis, L., Schroda, M., Stern, D., Umen, J., Willows, R., Wilson, N., Zimmer, S.L., Allmer, J., Balk, J., Bisova, K., Chen, C.J., Elias, M., Gendler, K., Hauser, C., Lamb, M.R., Ledford, H., Long, J.C., Minagawa, J., Page, M.D., Pan, J., Pootakham, W., Roje, S., Rose, A., Stahlberg, E., Terauchi, A.M., Yang, P., Ball, S., Bowler, C., Dieckmann, C.L., Gladyshev, V.N., Green, P., Jorgensen, R., Mayfield, S., Mueller-Roeber, B., Rajamani, S., Sayre, R.T., Brokstein, P., Dubchak, I., Goodstein, D., Hornick, L., Huang, Y.W., Jhaveri, J., Luo, Y., Martinez, D., Ngau, W.C., Otillar, B., Poliakov, A., Porter, A., Szajkowski, L., Werner, G., Zhou, K., Grigoriev, I.V., Rokhsar, D.S., Grossman, A.R., 2007. The Chlamydomonas genome reveals the evolution of key animal and plant functions. Science 318, 245–250. 10.1126/science.1143609

Mews, L.K., 1980. The Green Hydra Symbiosis. III. The Biotrophic transport of Carbohydrate from Alga to Animal. Proc. R. Soc. Lond., Ser. B: Biol. Sci. 209, 377–401. 10.1098/rspb.1980.0101

Mews, L.K., Smith, D.C., 1982. The Green Hydra Symbiosis. VI. What is the Role of Maltose Transfer from Alga to Animal? Proceedings of the Royal Society of London. Series B. Biological Sciences 216, 397–413. 10.1098/rspb.1982.0083

Moran, N.A., 2007. Symbiosis as an adaptive process and source of phenotypic complexity. Proc. Natl. Acad. Sci. U. S. A. 104, 8627–8633. 10.1073/pnas.0611659104

Murer, H., Biber, J., 1996. Molecular mechanisms of renal apical Na/phosphate cotransport. Annu. Rev. Physiol. 58, 607–618. 10.1146/annurev.ph.58.030196.003135

Muscatine, L., 1965. Symbiosis of hydra and algae. 3. Extracellular products of the algae. Comp. Biochem. Physiol. 16, 77–92

Muscatine, L., 1983. Hydra: Research Methods. Plenum Press, New York.

Muscatine, L., Lenhoff, H.M., 1963. Symbiosis: On the Role of Algae Symbiotic with Hydra. Science 142, 956–958. 10.1126/science.142.3594.956

Muscatine, L., Lenhoff, H.M., 1965a. Symbiosis of Hydra and Algae. I. Effects of Some Environmental Cations on Growth of Symbiotic and Aposymbiotic Hydra. Biol. Bull. 128, 415–424. 10.2307/1539903

Muscatine, L., Lenhoff, H.M., 1965b. Symbiosis of Hydra and Algae. II. Effects of Limited Food and Starvation on Growth of Symbiotic and Aposymbiotic Hydra. Biol. Bull. 129, 316–328

Muscatine, L., McAuley, P.J., 1982. Transmission of symbiotic algae to eggs of green hydra. Cytobios 33, 111–124

Nichols, H.W., Bold, H.C., 1965. Trichosacinapolymorpha Gen. et sp. NOV. Journal of Phycology 1, 34–38

Palenik, B., Grimwood, J., Aerts, A., Rouze, P., Salamov, A., Putnam, N., Dupont, C., Jorgensen, R., Derelle, E., Rombauts, S., Zhou, K., Otillar, R., Merchant, S.S., Podell, S., Gaasterland, T., Napoli, C., Gendler, K., Manuell, A., Tai, V., Vallon, O., Piganeau, G., Jancek, S., Heijde, M., Jabbari, K., Bowler, C., Lohr, M., Robbens, S., Werner, G., Dubchak, I., Pazour, G.J., Ren, Q., Paulsen, I., Delwiche, C., Schmutz, J., Rokhsar, D., Van de Peer, Y., Moreau, H., Grigoriev, I.V., 2007. The tiny eukaryote Ostreococcus provides genomic insights into the paradox of plankton speciation. Proc. Natl. Acad. Sci. U. S. A. 104, 7705–7710. 10.1073/pnas.0611046104

Pardy, R.L., 1976. The morphology of green hydra endosymbionts as influenced by host strain and host environment. J. Cell Sci. 20

Parra, G., Bradnam, K., Korf, I., 2007. CEGMA: a pipeline to accurately annotate core genes in eukaryotic genomes. Bioinformatics 23. 10.1093/bioinformatics/btm071

Perriere, G., Gouy, M., 1996. WWW-query: an on-line retrieval system for biological sequence banks. Biochimie 78, 364–369

Pombert, J.F., Blouin, N.A., Lane, C., Boucias, D., Keeling, P.J., 2014. A lack of parasitic reduction in the obligate parasitic green alga Helicosporidium. PLoS Genet. 10, e1004355. 10.1371/journal.pgen.1004355

Prochnik, S.E., Umen, J., Nedelcu, A.M., Hallmann, A., Miller, S.M., Nishii, I., Ferris, P., Kuo, A., Mitros, T., Fritz-Laylin, L.K., Hellsten, U., Chapman, J., Simakov, O., Rensing, S.A., Terry, A., Pangilinan, J., Kapitonov, V., Jurka, J., Salamov, A., Shapiro, H., Schmutz, J., Grimwood, J., Lindquist, E., Lucas, S., Grigoriev, I.V., Schmitt, R., Kirk, D., Rokhsar, D.S., 2010. Genomic analysis of organismal complexity in the multicellular green alga Volvox carteri. Science 329, 223–226. 10.1126/science.1188800

Quesada, A., Galvan, A., Fernandez, E., 1994. Identification of nitrate transporter genes in Chlamydomonas reinhardtii. Plant J. 5, 407–419

Rahat, M., Reich, V., 1984. Intracellular infection of aposymbiotic Hydra viridis by a foreign free-living Chlorella sp.: initiation of a stable symbiosis. J. Cell Sci. 65, 265–277

Rees, T.A.V., 1986. The Green Hydra Symbiosis and Ammonium I. The Role of the Host in Ammonium Assimilation and its Possible Regulatory Significance. Proc. R. Soc. Lond., Ser. B: Biol. Sci. 229, 299–314. 10.1098/rspb.1986.0087

Rees, T.A.V., 1989. The Green Hydra Symbiosis and Ammonium II. Ammonium Assimilation and Release by Freshly Isolated Symbionts and Cultured Algae. Proc. R. Soc. Lond., Ser. B: Biol. Sci. 235

Rees, T.A.V., 1991. Are Symbiotic Algae Nutrient Deficient? Proc. R. Soc. Lond. B. Biol. Sci. 243, 227–233. 10.1098/rspb.1991.0036

Roffman, B., Lenhoff, H.M., 1969. Formation of Polysaccharides by Hydra from Substrates produced by their Endosymbiotic Algae. Nature 221, 381–382

Sanz-Luque, E., Chamizo-Ampudia, A., Llamas, A., Galvan, A., Fernandez, E., 2015. Understanding nitrate assimilation and its regulation in microalgae. Front Plant Sci 6, 899. 10.3389/fpls.2015.00899

Schwentner, M., Bosch, T.C., 2015. Revisiting the age, evolutionary history and species level diversity of the genus Hydra (Cnidaria: Hydrozoa). Mol. Phylogenet. Evol. 91, 41–55. 10.1016/j.ympev.2015.05.013

Shinzato, C., Shoguchi, E., Kawashima, T., Hamada, M., Hisata, K., Tanaka, M., Fujie, M., Fujiwara, M., Koyanagi, R., Ikuta, T., Fujiyama, A., Miller, D.J., Satoh, N., 2011. Using the Acropora digitifera genome to understand coral responses to environmental change. Nature 476, 320–323. 10.1038/nature10249

Stanke, M., Keller, O., Gunduz, I., Hayes, A., Waack, S., Morgenstern, B., 2006. AUGUSTUS: ab initio prediction of alternative transcripts. Nucleic Acids Res. 34. 10.1093/nar/gkl200

Tao, T.Y., Towle, H.C., 1986. Coordinate regulation of rat liver genes by thyroid hormone and dietary carbohydrate. Ann. N. Y. Acad. Sci. 478, 20–30

Tautz, D., Domazet-Loso, T., 2011. The evolutionary origin of orphan genes. Nat. Rev. Genet. 12, 692–702. 10.1038/nrg3053

The Gene Ontology, C., Ashburner, M., Ball, C.A., Blake, J.A., Botstein, D., Butler, H., Cherry, J.M., Davis, A.P., Dolinski, K., Dwight, S.S., Eppig, J.T., Harris, M.A., Hill, D.P., Issel-Tarver, L., Kasarskis, A., Lewis, S., Matese, J.C., Richardson, J.E., Ringwald, M., Rubin, G.M., Sherlock, G., 2000. Gene Ontology: tool for the unification of biology. Nat. Genet. 25, 25–29. 10.1038/75556

Thorington, G., Margulis, L., 1981. Hydra viridis: transfer of metabolites between Hydra and symbiotic algae. The Biological Bulletin 160, 175–188. 10.2307/1540911

Vandermeulen, J.H., Davis, N.D., Muscatine, L., 1972. The effect of inhibitors of photosynthesis on zooxanthellae in corals and other marine invertebrates. Mar. Biol. 16, 185–191. 10.1007/bf00346940

Worden, A.Z., Lee, J.H., Mock, T., Rouze, P., Simmons, M.P., Aerts, A.L., Allen, A.E., Cuvelier, M.L., Derelle, E., Everett, M.V., Foulon, E., Grimwood, J., Gundlach, H., Henrissat, B., Napoli, C., McDonald, S.M., Parker, M.S., Rombauts, S., Salamov, A., Von Dassow, P., Badger, J.H., Coutinho, P.M., Demir, E., Dubchak, I., Gentemann, C., Eikrem, W., Gready, J.E., John, U., Lanier, W., Lindquist, E.A., Lucas, S., Mayer, K.F., Moreau, H., Not, F., Otillar, R., Panaud, O., Pangilinan, J., Paulsen, I., Piegu, B., Poliakov, A., Robbens, S., Schmutz, J., Toulza, E., Wyss, T., Zelensky, A., Zhou, K., Armbrust, E.V., Bhattacharya, D., Goodenough, U.W., Van de Peer, Y., Grigoriev, I.V., 2009. Green evolution and dynamic adaptations revealed by genomes of the marine picoeukaryotes Micromonas. Science 324, 268–272. 10.1126/science.1167222

